# Mito-SinCe^2^ Approach to Quantify Relationships between Mitochondrial Dynamics and Energetics status

**DOI:** 10.1101/537373

**Authors:** Brian Spurlock, Priyanka Gupta, Malay Kumar Basu, Avik Mukherjee, Anita B Hjelmeland, Victor Darley-Usmar, Danitra Parker, McKenzie E Foxall, Kasturi Mitra

## Abstract

Mitochondrial dynamics primarily refers to the opposing fission and fusion events between individual mitochondria, which maintains steady-state mitochondrial morphology in cells. The details of the bidirectional relationship between mitochondrial dynamics and energetics are unclear as the methods to integrate these mitochondrial aspects are lacking. To study the quantitative relationship between mitochondrial dynamics (fission, fusion, matrix continuity and diameter) and energetics (ATP and redox), we have developed and validated an analytical approach, mito-SinCe^2^. Application of this approach led to the hypothesis that mitochondria-dependent ovarian tumor initiating cells interconvert between 3 states with distinct relationships between mitochondrial energetics and dynamics. Interestingly, fusion and ATP increase linearly with each other only once a certain level of fusion is attained. Moreover, mitochondrial dynamics status changes linearly either with ATP or with redox, but not simultaneously with both. Furthermore, mito-SinCe^2^ analyses can potentially predict new quantitative features of the opposing fission vs. fusion relationship and classify cells into functional classes based on their mito-SinCe^2^ states.

## Introduction

The steady-state morphology of mitochondria within a given cell depends on the cell type and its physiological state (Miettinen and Bjorklund, 2017). Mitochondrial morphology is primarily determined by a balance of opposing fission and fusion events between individual mitochondria, which are collectively referred to as mitochondrial dynamics (Nunnari and Suomalainen, 2012). Mitochondrial dynamics (opposing fission and fusion) and mitochondrial energetics (linked ATP and redox states (Murphy, 2009)) impact each other to regulate cellular bioenergetics (Mishra and Chan, 2016; Willems et al., 2015). In nutrient excess the balance of mitochondrial dynamics lies towards fission, decreasing mitochondrial energetic efficiency. Conversely, in nutrient deprivation, mitochondria are maintained in hyperfused networks, enhancing mitochondrial energetic efficiency (Liesa and Shirihai, 2013; Schrepfer and Scorrano, 2016).

In-depth understanding of the relationship of mitochondrial dynamics and energetics or its underlying mechanisms require integrated quantitative analyses of these mitochondrial properties. Various approaches to quantify mitochondrial energetics exist in the field. Quantification of mitochondrial morphology reflecting mitochondrial dynamics is less developed, but has been attempted by various laboratories (Harwig et al., 2018; Iannetti et al., 2016; Karbowski et al., 2004; Mitra et al., 2009; Ouellet et al., 2017; Parker et al., 2017). To address the critical need for integrated analyses, we first designed and validated metrics for quantifying mitochondrial dynamics status and thereafter developed a multivariate approach towards identifying and quantifying mitochondrial [energetics] vs. [dynamics] relationships in single cells. We named the microscopy based high resolution approach as mito-SinCe^2^ (mito-SinCe-SQuARED: <u>Sin</u>gle <u>Ce</u>ll <u>S</u>imultaneous <u>Qu</u>antification of <u>A</u>TP or <u>Re</u>dox with <u>D</u>ynamics of <u>Mito</u>chondria). After validating and providing proof of principal of the mito-SinCe^2^ approach, we used it in an example application on stem cell regulation.

The importance of mitochondrial dynamics and energetics in stem cell regulation is becoming increasingly recognized (Chen and Chan, 2017). Previously, we proposed that fission activity is critical for ovarian cancer (Tanwar et al., 2016). Here we find that mitochondria dependent ovarian tumor initiating cells (ovTICs), which harbor stem cell properties and contribute to ovarian cancer development (Ishiguro et al., 2016), regulate mitochondrial energetics differently than the bulk tumor cells. Mito-SinCe^2^ analyses reveal complexities of mitochondrial dynamics and energetics in ovTIC self-renewal and proliferation.

## Results

### Development and validation of single cell metrics for quantifying the state of mitochondrial dynamics

Single cell metrics for parsing the contribution and dynamic interaction of the opposing fission or fusion processes in a given mitochondrial morphology have not been reported. Here, we used outputs of MitoGraph v2.1 image analysis software (Viana et al., 2015) and photoactivation approach to compute and validate the quantitative metrics for steady state mitochondrial fission and fusion and also average mitochondrial diameter in single cells.

Mitograph v2.1 application on high resolution 3-D confocal micrographs yields the following parameters for mitochondrial elements: total length and number, individual length and volume. First, we confirmed that the length and diameter (derived from volume) of mitochondrial elements correspond with qualitative differences in mitochondrial morphology in representative Mitotracker Green (MTG) stained cells 1-4 (**Fig. 1A**, details in Suppl Results, **Fig. S1A,B**). Mitochondrial fission primarily (along with biogenesis and clearance) determines the number of mitochondrial elements in a cell (Liesa and Shirihai, 2013). Thus, we obtained a quantitative fission metric, denoted [Fission], by normalizing the total number of mitochondrial elements (N) within a cell by the total length of its mitochondrial network (ΣL) (to account for variations in total mitochondrial content). On the other hand, mitochondrial fusion may increase the length of individual mitochondrial elements (Mitra et al., 2009). Thus, we obtained quantitative fusion metrics, denoted [Fusion(1-10)], from the percent coverage of the top(1-10) longest elements. We obtained [Fusion(1,3,5,10)] by normalizing the longest (L_1_), or the summed length of three (L_3_), or five (L_5_) or ten (L_5_) longest components by the total length of the mitochondrial network (ΣL). Therefore, [Fission]_(N / ΣL)_ relies on mitochondrial number and [Fusion]_(L1-10 / ΣL)_ relies on mitochondrial length, and thus report distinct structural properties of a mitochondrial network. We also calculated the mean mitochondrial diameter in each cell, denoted [Diameter], by weighting the diameter of each element by its volume (details in Suppl. results). We found that Cells 1 and 2 have higher [Fusion(1-10)] and lower [Fission] in comparison to Cells 3 and 4, consistent with the qualitative evaluation of the images (**Fig. 1A, B**). The higher [Diameter] of Cell 3 is due to several small and thick mitochondria, whereas the non-uniformly thick mitochondrial tubules of Cell 1 do not increase [Diameter] (other minor limitation in Suppl. results) (**Fig. 1A, B**).

**Fig. 1.**
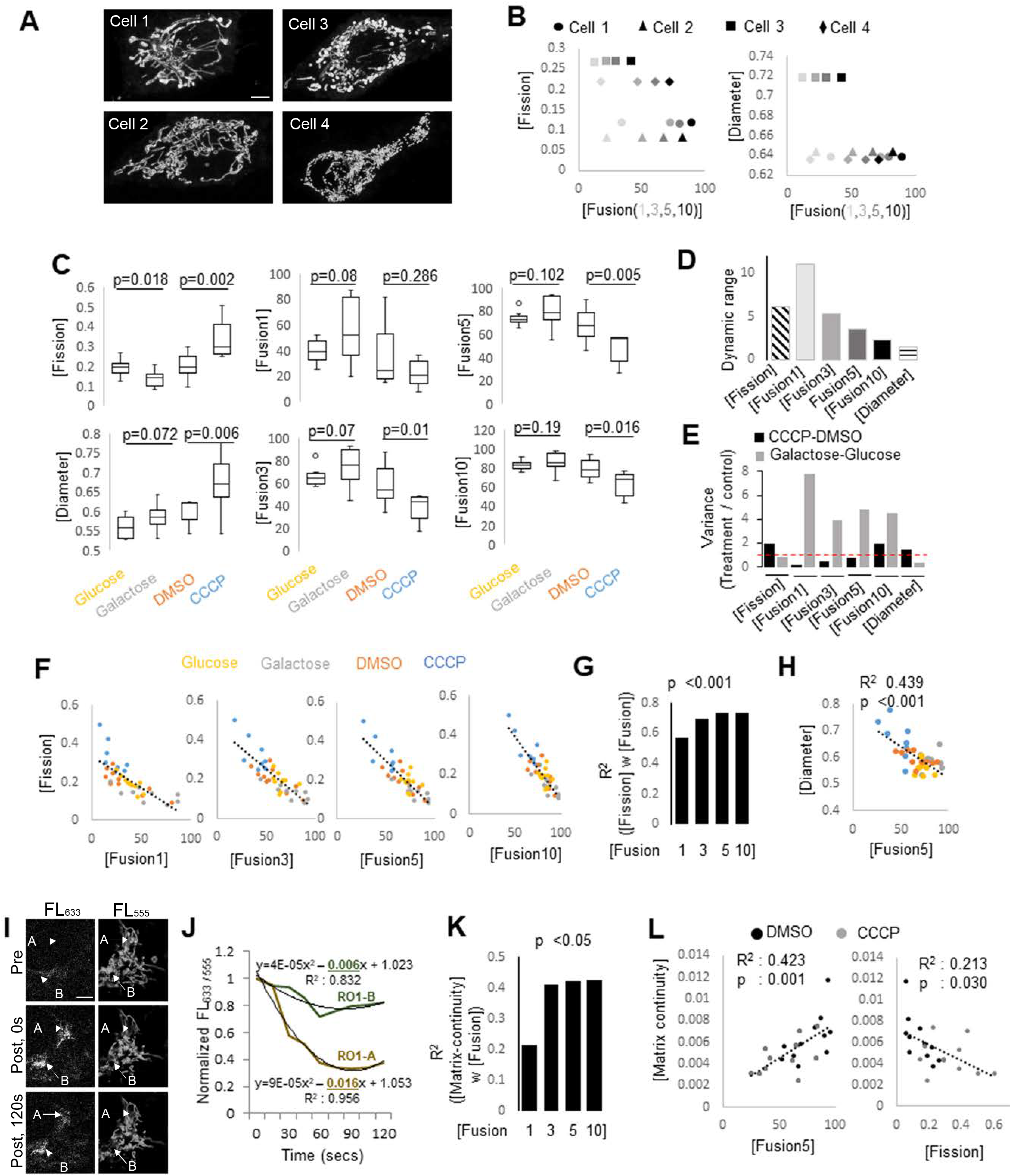
Development and validation of single cell metrics to quantify the state of mitochondrial dynamics. **(A)** Confocal micrographs of A2780-CPs representing different mitochondrial morphology reflecting state of dynamics in cells 1-4. Scale bar is 5 µm. **(B)** Bivariate plot of [Fission] or [Diameter] with [Fusion(1, 3, 5,10)] from cells 1-4 in A. **(C)** Box plot showing [Fission], [Fusion1, 3, 5, 10] and [Diameter] in HaCaT cells maintained in fresh Glucose, Galactose, DMSO or CCCP for 2 hrs. p values are from Kruskal-Wallis test. **(D)** Bar plot showing dynamic range obtained from C. **(E)** Bar plot of variance of metrics in C. Dashed line represents no difference between treatment and control. **(F)** Bivariate plot for linear regression analyses of [Fission] with [Fusion(1,3,5,10)] from experiment described in C. Regression lines shown. **(G)** Bar plot of R^2^ values of the regression lines from C, while p value is shown. **(H)** Bivariate plot for linear regression analyses of [Diameter] with [Fusion5] from experiment described in C. Regression lines with R^2^ and p values shown. **(I)** Representative confocal micrographs of a mito-PSmO2 expressing HaCaT cell, depicting photo-switching based matrix continuity assay. A and B are ROIs photo-switched and monitored over time (arrows). Scale bar is 5 µm. **(J)** Quantitation of fluorescence signal from ROIs A and B in I. See methods for details. **(K)** Bar plot of the R^2^ values of linear regression analyses between [Matrix-continuity] and [Fusion(1,3,5,10)] of the same single HaCaT cells, while p value is shown. **(L)** Bivariate plot for linear regression analyses of [Fission] or [Fusion5] with [Matrix-continuity] of the HaCaT cells in K.

The first level of validation of our metrics for fission or fusion involved 2 hrs exposure to galactose or CCCP, since the protonophore CCCP prevents and galactose promotes fusion, thus altering the steady state mitochondrial morphology (Mishra and Chan, 2016). We used immortalized human cells (HaCaT) stably expressing our mitochondrial-matrix targeted Photo-Switchable mOrange2 probe (mito-PSmO2), whose basal fluorescence can be photo-switched (Subach et al., 2011). Using the basal mito-PSmO2 fluorescence we detected expected differences in fission and fusion when comparing CCCP with DMSO, and galactose with glucose (**Fig. 1C**). We detected higher [Fission] and lower [Fusion(1-10)] in CCCP treated cells, with opposite trends in cells in galactose. Also, CCCP treated cells have higher [Diameter]. The dynamic range (Value_max_ / Value_min_) is maximal for [Fusion1] and progressively decreases with [Fusion(1-10)] (**Fig. 1D**), indicating that inclusion of fewer elements in [Fusion1] increases metric sensitivity. We reasoned that enhancement or suppression of any biological process would respectively increase or decrease the variability in the metric measuring the process. Indeed, the variation (expressed as variance) in [Fusion(1-10)], and not [Fission], is markedly reduced with CCCP treatment that suppresses fusion and increased in galactose that enhances fusion (**Fig. 1E**). With linear regression analyses of the single cell data (assessed by R^2^ and p value), we found an inverse [Fission] vs. [Fusion(1-10)] relationship i.e. [Fission] increases linearly with decrease in [Fusion(1-10)] **(Fig. 1F)**. Optimal linearity is achieved with Fusion5 or above (R^2^ and p in **Fig. 1G**), indicating that inclusion of more long elements (5 or 10) better represent the overall fusion status of the cell. This data validates the consensus that fission and fusion counter each other (Hoppins, 2014) and reveals the quantitative relationship between opposing fission and fusion processes contributing to steady state mitochondrial morphology. We also find [Diameter] decreases linearly with increase in [Fusion(1-10)] (**Fig. 1H** shows only Fusion[5]), the significance of which remains unknown.

To further validate [Fusion(1-10)] obtained from MitoGraph v2.1 analysis of diffraction limited micrographs, we assessed matrix continuity arising due to fusion of the double membranes between contiguous mitochondria. We refined our previously reported matrix-continuity assay (Mitra et al., 2009) by photo-switching mito-PSmO2 and simultaneously measuring the dilution of the photo-switched fluorescence (FL_633_) and the recovery of the bleached basal fluorescence (FL_555_) (**Fig. 1I**); distinction of the matrix-continuity assay from reported fusion assays is in Suppl. Method (**Fig. S1C**). We denoted the linear coefficient of the fitted curve for decay of FL_Ex-633/555_ in a region of interest (ROI) as its [Matrix-continuity] (**Fig. 1J**). As expected, [Matrix-continuity] is less in the CCCP treated cells (**Fig. S1D**). Importantly, linear regression analyses indicates matrix continuity increases linearly with fusion and thus validates [Fusion(1-10)] (**Fig. 1K, L**). Notably, [Matrix-continuity] decreases linearly with [Fission], but only modestly. The dynamic range of [Fusion5] is comparable to that of [Matrix-continuity], while that of [Fusion1] remains maximum (**Fig. S1E**).

Finally, we validated [Fission] and [Fusion(1,5)] using Mitotracker-633 stained mouse embryonic fibroblasts (MEFs) lacking the key fusion proteins MFN-1/2 (Chen et al., 2003), or the key fission protein Drp1 (Ishihara et al., 2009). As expected, MFN-1/2 DKO exhibits higher [Fission] and lower [Fusion5] as compared to its corresponding wild type, WTm (**Fig. 2A,B**). Also, MFN-1/2 DKO exhibits a robust increase in [Diameter], as validated by direct manual measurements (in μm) (**Fig. S1F**). Consistent with the loss of regulation of fusion in the absence of MFN1/2, the variance of [Fusion(1,5)] is markedly reduced in MFN1/2-DKO in comparison to WTm (**Fig. 2C**). As expected, the inverse [Fission or Diameter] vs. [Fusion5] linear relationship present in WTm is lost in MFN1/2-DKO (**Fig. 2D**). Similar comparison of Drp1-KO and its corresponding wild type MEFs, WTd, shows significant changes only in [Diameter] (**Fig. 2E, F)**. However, maximal reduction of the variance of [Fission] in Drp1-KO in comparison to WTd (**Fig. 2G**) is consistent with loss of regulation of fission by Drp1. The inverse [Fission] vs. [Fusion5] linear relationship is modestly modified in Drp1-KO. (**Fig. 2H**, left, reflected in modest change in slope of regression lines). The inverse [Diameter] vs. [Fusion5] relationship is not affected (**Fig. 1H**, right), while [Diameter] is markedly elevated in Drp1-KO (**Fig. 2H,** right).

**Fig. 2.**
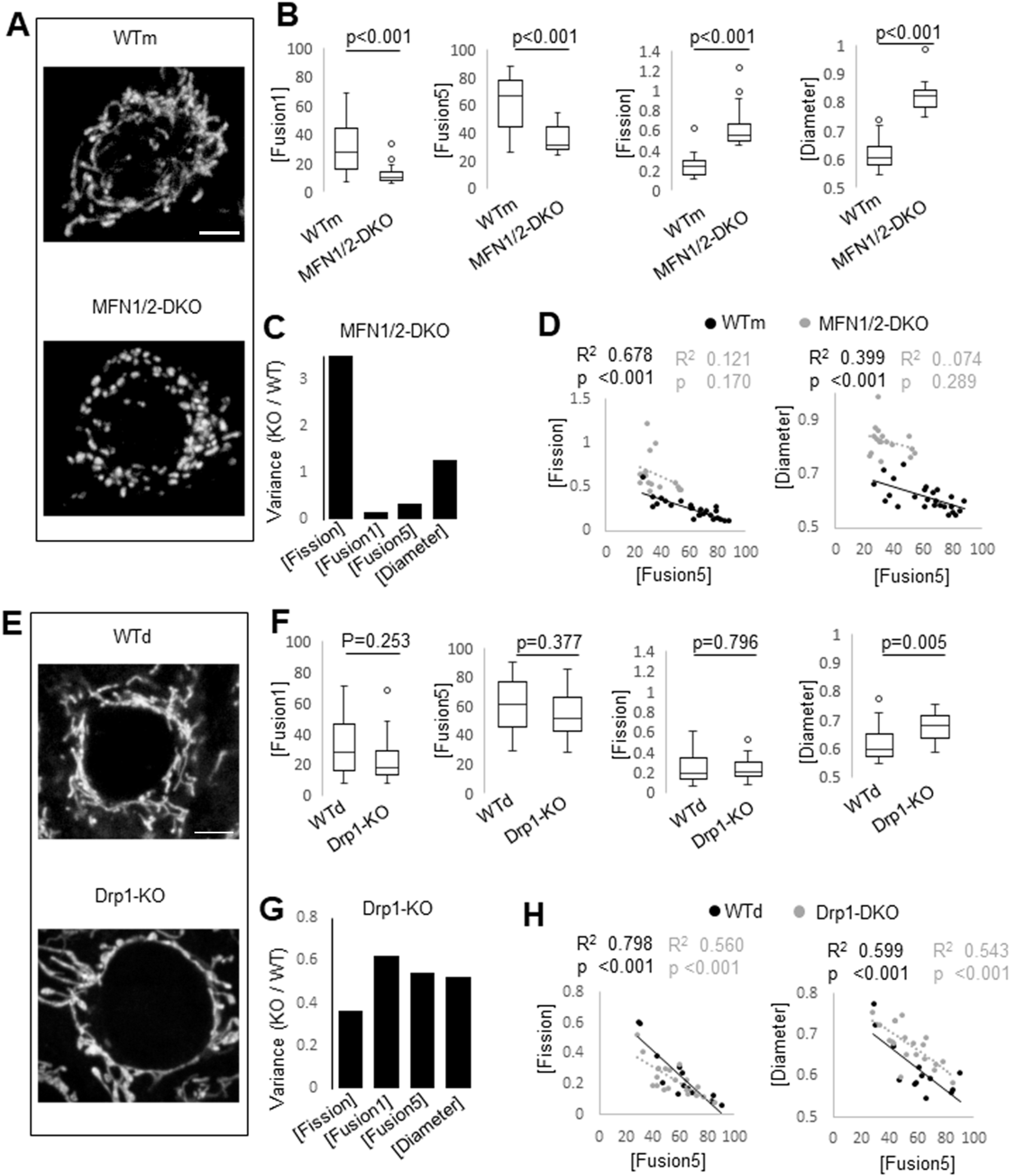
Validation of dynamics metrics in Drp1 and MFN1/2 ablated cells. **(A)** Representative confocal micrographs of MFN1/2-DKO and WTm MEFs. Scale bar is 5 µm. **(B)** Box plot showing [Fusion(1,5)], [Fission] and [Diameter] in WTm and MFN1/2-DKO MEFs. p values are from Kruskal-Wallis test. **(C)** Bar plots of variance of metrics from B. **(D)** Bivariate plot for linear regression analyses of [Fission] or [Diameter] with [Fusion-5] in WTm and MFN1/2-DKO MEFs. Regression lines with R^2^ and p values shown. **(E)** Representative confocal micrographs of Drp1-KO and WTd MEFs. Scale bar is 5 µm. **(F)** Box plot showing [Fusion(1,5)], [Fission] and [Diameter] in WTd and Drp1-KO MEFs. p values are from Kruskal-Wallis test. **(G)** Bar plots of variance of metrics from E. **(H)** Bivariate plot for linear regression analyses of [Fission] or [Diameter] with [Fusion-5] in WTd and Drp1-KO MEFs. Regression lines with R^2^ and p values shown.

In summary, using various mitochondrial structural probes, we identified quantitative metrics for opposing fission / fusion processes that contribute to mitochondrial dynamics state in single cells; these steady state metrics do not quantify fission or fusion kinetics (minor limitations in Suppl. Info.). We validated the metrics in metabolically distinct cells and in fission / fusion mutants.

### Development and validation of mito-SinCe^2^ approach towards quantifying the relationships between the status of mitochondrial dynamics and energetics

To identify and quantify mitochondrial [energetics] vs. [dynamics] relationships, we designed the high resolution confocal microscopy based approach, mito-SinCe^2^. This involves single cell quantitative analyses of the state of mitochondrial dynamics (using metrics for fission, fusion, matrix continuity and diameter as in **Fig. 1, 2**) and energetics (metrics for ATP or redox, using genetically encoded fluorescent ratiometric probes to rule out influence of mitochondrial mass). The mito-SinCe^2^ workflow involves two independent arms: Dynamics-ATP and Dynamics-Redox (**Fig. 3A**, **Fig. S1G,H**). The ATP arm uses the mito-GO-ATeam2 FRET probe (FL_Em-555/488_) (Nakano et al., 2011) to quantify the basal steady state ATP levels in the mitochondrial matrix, denoted [ATP]. We used the direct excitation of the FRET acceptor (FL_Ex-555_) to measure dynamics. The Redox arm uses the mito-roGFP probe (FL_Ex-405/488_) to quantify the oxidation status of the mitochondrial matrix(Hanson et al., 2004), denoted [Oxidation]. The dynamics metrics obtained with the functional mito-roGFP probe are not different between the CCCP and galactose treated cells (**Fig. S1I**), indicating mito-roGFP cannot be used as a structural probe. Therefore, we used the fluorescently compatible mito-PSmO2 probe to quantify dynamics, in combination with mito-roGFP (as validated in **Fig. 1F, G)**.

**Fig. 3.**
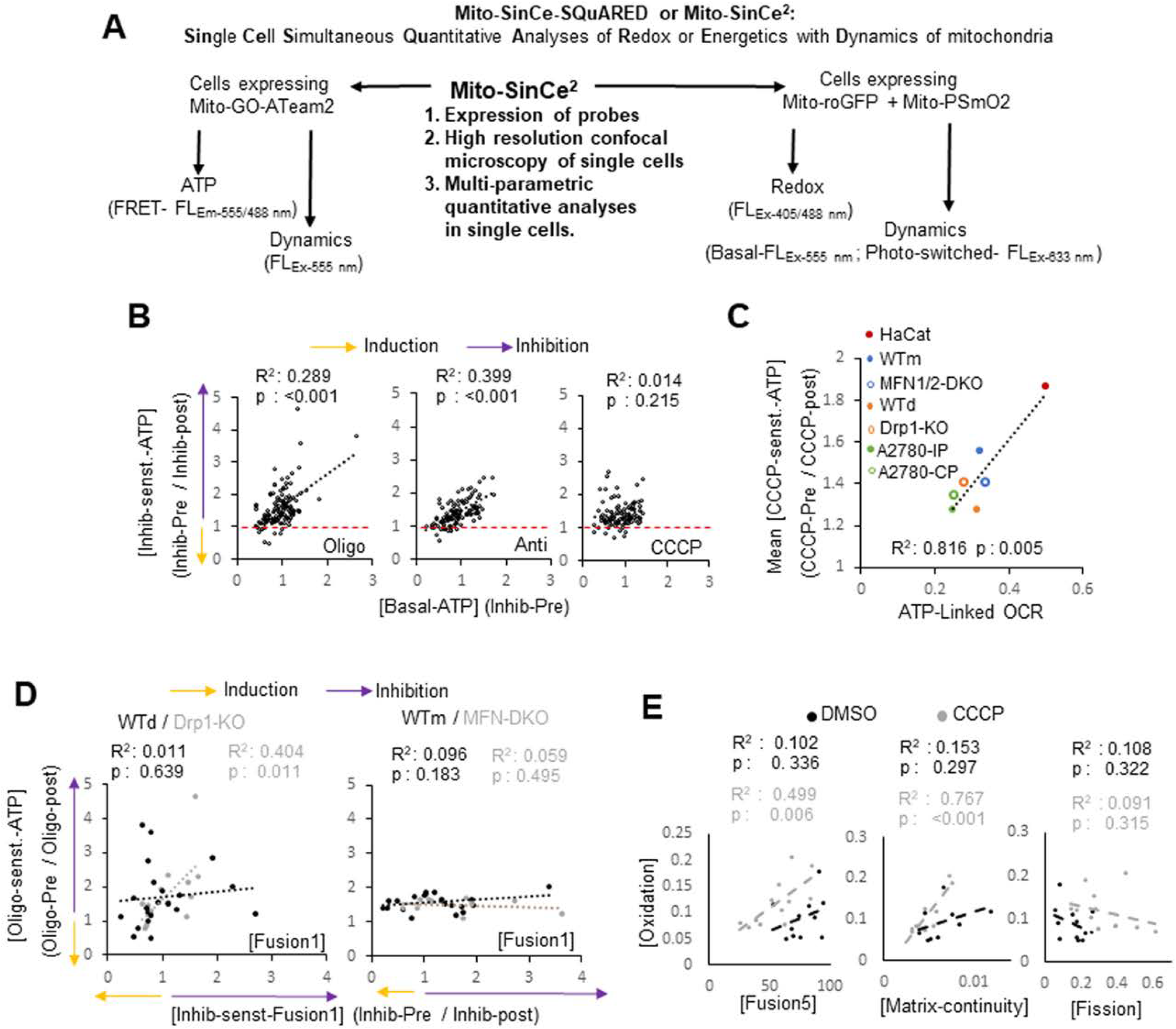
Development and validation of the mitoSinCe^2^ approach. **(A)** Schematic of the mito-SinCe^2^ work flow. **(B)** Bivariate plot for regression analyses of [Oligomycin or Antimycin or CCCP-sensitive-ATP] and [Basal-ATP] in 7 cell lines as detected by the mito-GO-ATeam reporter. Regression lines with R^2^ and p values shown. Dashed line denoting no induction or inhibition. **(C)** Bivariate plot for linear regression analyses of the mean [CCCP-sensitive-ATP] for each cell line and ATP-linked-OCR. Regression lines with R^2^ and p values shown. **(D)** Mito-SinCe^2^ (ATP-arm) analyses [Oligo-sensitive-ATP] and [Oligo-sensitive-Fusion1] in Drp1KO / Wtd and Drp1-KO / WTm cell pairs. Regression lines with R^2^ and p values shown. **(E)** Mito-SinCe^2^ (redox-arm) analyses of [Oxidation] with [Fusion5], [Matrix-continuity] and [Fission] in the presence of CCCP or DMSO in HaCaTs.

First, we validated the mito-roGFP and mito-GoATeam2 probes. For the widely used mito-roGFP probe, we confirmed that oxidizing agent t-BH increases [Oxidation] and reducing agent DTT causes no reduction beyond the basal redox levels (**Fig. S1J**). It is noteworthy that mito-roGFP detects oxidation status but does not detect reactive oxygen species directly. We extensively validated the newer mito-GO-ATeam2 probe for quantification of both ATP and dynamics. Mitochondrial ATP synthesis is coupled to oxygen consumption via the transmembrane potential (ΔΨ) maintained by electron transport chain complexes (Hill et al., 2012). To validate mito-GO-ATeam2 in 7 cell lines, we used mitochondrial inhibitors that reduce mitochondrial ATP levels. The inhibitors used are Oligomycin (inhibits ATP-synthase/ATPase), Antimycin (inhibits complex III) and CCCP (uncouples ATP synthesis and oxygen consumption by dissipating ΔΨ) (Brand and Nicholls, 2011). The cell lines used are paired WTd and Drp1-KO and WTm and MFN1/2-DKO MEFs, normal human HaCaT line, and paired A2780-IP and A2780-CP ovarian cancer lines. We excluded the cells (∼10%) where mito-GO-ATeam2 fluorescence is enriched in mitochondrial specks of unknown significance (basic characterization in Suppl. results). Here, we exposed cells to the inhibitors (10 mins) and computed the Pre-Inhibitor / Post-Inhibitor ratio of [ATP] to signify the [Inhibitor-sensitive-ATP], while the ‘Pre’ value signifies the [Basal-ATP] (**Fig. 3B**); [Inhibitor-sensitive-ATP] value >1 indicates inhibition, while 1 indicates no change. Using mito-GO-ATeam2, we detected 1 to >4 fold reduction of [Basal-ATP] caused by the mitochondrial inhibitors. Under the same conditions these inhibitors impact Oxygen Consumption Rate (OCR) measured by Seahorse metabolic flux analyses (**Fig. S1K**). [Oligomycin- or Antimycin-sensitive-ATP] increases with increase in [Basal-ATP], which is not observed for [CCCP-sensitive-ATP] (**Fig. 3B**). Importantly, as expected, the mean [CCCP-sensitive-ATP] for each cell line, obtained using mito-GO-ATeam2, increases with increase in the [ATP-linked-OCR] of that cell line, obtained using metabolic flux analyses (**Fig. 3C**). Thus, we could successfully validate the use of mito-GO-ATeam2 as an ATP probe in mito-SinCe^2^. The use of mito-GO-ATeam2 for measuring the metrics for dynamics was validated by the following results: **a)** inverse [Fission] vs. [Fusion5] relationship (**Fig. S1L**); **b)** CCCP (10 mins) induced decrease in [Fusion (1,5)] in 6 lines but not consistently in MFN1/2-DKO (**Fig. S1M**); **c)** expected reduction in [Fission] in the Drp1-KO and decrease in [Fusion (1,5)] in the MFN1/2-DKO in comparison to their respective WT MEFs (not shown). We found Mito-GO-ATeam2 is more sensitive than Mitotracker-633 in detecting difference between WT and KO MEFs (other differences between probes are in limitations in Suppl info). Notably, CCCP, which reduced fusion (Mishra and Chan, 2016), selectively reduces variance of [Fusion(1,5)] combined from all cell lines (**Fig. S1N**), similar to that of fusion defective MFN1/2-DKO cells (**Fig. 2C**).

Next, we provide proof of principal of the ATP arm of mito-SinCe^2^. Unopposed mitochondrial fusion has been linked to ATP output. Therefore, we used the ATP arm of the mito-SinCe^2^ to analyze the impact of Oligomycin induced inhibition of mitochondrial ATP synthase on the dynamics metrics, in the presence and absence of the key fission and fusion proteins Drp1 and MFN1/2 respectively. Thus, we performed linear regression analyses of Oligomycin sensitive [ATP] and [Fusion] (experimental plan as in Fig. 3B); > 1 value indicates inhibition, 1 indicates no change, and < 1 indicates induction (**Fig. 3D**). [Oligomycin-sensitive-Fusion1] increases linearly with [Oligomycin-sensitive-ATP] in the Drp1-KO MEFs, while this linear relationship was not significant in the WTd MEFs (**Fig. 3D,** left). A similar trend was observed with [Oligomycin-sensitive-Fusion5] (not shown). This indicates that fusion decreases linearly with inhibition of ATP synthesis in the absence of Drp1, while any non-linear nature of the relationship in the presence of Drp1 remains to be identified. Thus, our data indicates that Drp1 may modify the reported regulation of fusion by ATP (Mishra and Chan, 2016). No statistically significant linear relationship was observed in the MFN-1/2-DKO / WTm pair (**Fig. 3D,** right). Note that the WTd MEFs, obtained at embryonic stage E10 (Ishihara et al., 2009), and WTm MEFs, obtained at embryonic stage E13 (Chen et al., 2003), have different spreads of data in the bivariate plots.

Next, we provide proof of principal of the redox arm of mito-SinCe^2^. Here, we investigated the impact of CCCP driven disruption of mitochondrial potential, given CCCP can either induce or prevent mitochondrial oxidation (Izeradjene et al., 2005; Murphy, 2009). We found that [Oxidation] linearly increases with [Fusion5] and [Matrix-continuity] within the CCCP treated population, while the linear relationship was not statistically significant in DMSO treated cell population (**Fig. 3E**, left and middle). Notably, CCCP increases [Oxidation] only in cells having [Fusion5] comparable to that of DMSO (**Fig. 3E,** left). It remains to be tested if the CCCP driven oxidation in cells with higher mitochondrial fusion is due to elevated oxidative in these cells. No statistically significant linear relationship was detected between [Oxidation] and [Fission] (**Fig. 3E,** right).

Thus, we provide proof of principal of the mito-SinCe^2^ approach by validating and refining reported findings. Hereafter, we will employ the approach in addressing biologically relevant questions (**Fig. 4-7**), and generate predictions for future experimental validations (**Fig. 9**). To investigate the various other aspects of quantitative relationships between the status of mitochondrial dynamics and ATP or redox, the data generated by mito-SinCe^2^ can be meaningfully studied in various other ways supported by appropriate statistics.

**Fig. 4.**
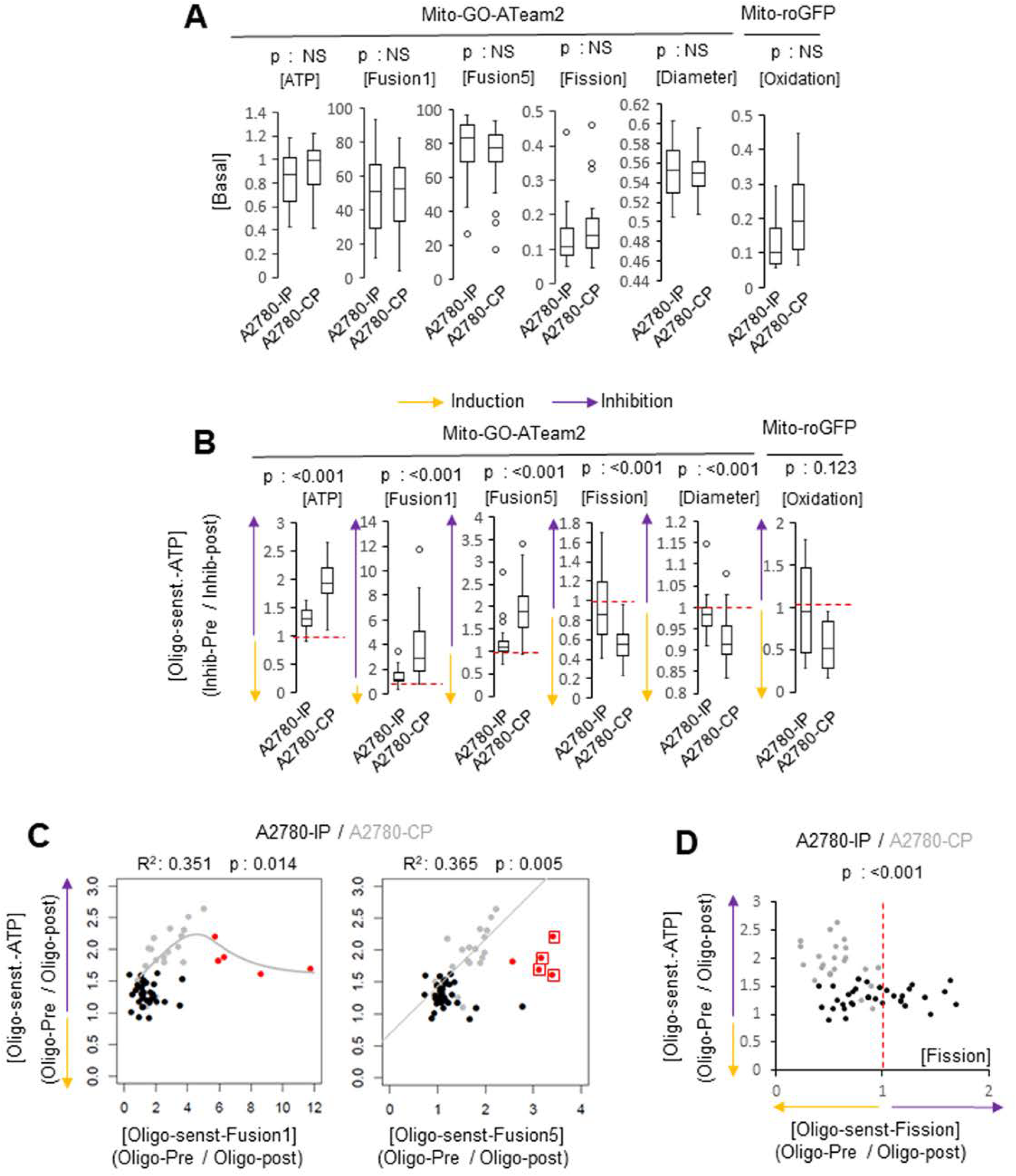
MitoSinCe^2^ comparison of the chemosensitive A2780-IPs and chemoresistant A2780-CPs. **(A)** Box plots of basal energetics and dynamics metrics. p values are from Kruskal-Wallis test. **(B)** Box plots of Oligomycin-sensitive energetics and dynamics metrics. Dashed line denoting no induction or inhibition. p values are from Kruskal-Wallis test. **(C)** Bivariate plot for non-linear regression analyses of [Oligo-sensitive-ATP] and [Oligo-sensitive-Fusion (1, 5)]. Regression lines with R^2^ and p values shown. **(D)** Bivariate plot analyses for Fisher’s Exact analyses (two-sided) of [Oligo-sensitive-ATP] and [Oligo-sensitive-Fission]. Dashed line denoting no induction or inhibition.

### Mito-SinCe^2^ analyses reveals unique relationships between status of mitochondrial dynamics and energetics in ovTIC^Aldh+^ enriched ovarian cancer cells

Given our interest in ovarian cancer (Tanwar et al., 2016), we compared the chemosensitive A2780-IP and the chemoresistant A2780-CP cells derived after carboplatin treatment (Landen et al., 2010) that enriches ovarian tumor initiating cells (ovTICs) (Ishiguro et al., 2016). Thus, compared to A2780-IPs, A2780-CPs are enriched in cells with enhanced activity of a functional ovTIC marker, Aldehyde dehydrogenase (Aldh) (detected by ALDEFLUOR stain) (Landen et al., 2010)(**Fig. S2A**). Using, mito-SinCe^2^ on A2780-IPs and A2780-CPs, we did not find any difference in basal [ATP], [Fission], [Fusion(1,5)], [Diameter] or [Oxidation] between the cell lines (**Fig. 4A**). However, compared to the A2780-IPs, the A2780-CPs have significantly higher [Oligomycin-sensitive-ATP] associated with greater Oligomycin induced decrease in [Fusion(1,5)], and increase in [Fission], [Diameter], while increase in [Oxidation] remained insignificant (**Fig. 4B**). These data demonstrate that steady state mitochondrial ATP levels and dynamics status are more dependent on active ATP synthesis in the ovTIC^Aldh+^ enriched A2780-CPs cells than the A2780-IPs. In contrast, mitochondrial energetics and dynamics status are equally dependent on mitochondrial transmembrane potential in the A2780-IPs and A2780-CPs, since the cell lines are comparably impacted by disruption of mitochondrial potential (by CCCP) or inhibition of mitochondrial complex III (by Antimycin) (**Fig. S2B,C**). Bivariate analyses of [Oligomycin-sensitive-ATP] and [Oligomycin-sensitive-Fusion1] distinguishes the A2780-CP cells contributing to the higher average [Oligomycin-sensitive-ATP] (**Fig. 4C**). Interestingly, [Oligomycin-sensitive-ATP] increases with [Oligomycin-sensitive-Fusion1] upto a certain level and decreases thereafter (as indicated by a nonlinear regression model) (**Fig. 4C**, left); no such relationship is observed in the A2780-IPs. Thus, specifically in the A2780-CPs, Oligomycin driven inhibition of ATP synthesis inhibits fusion only till a certain threshold (grey cells) beyond which this relationship does not hold true (red cells). These red cells are identified as distinct outliers (by interquartile range analyses) in the bivariate analyses of [Oligomycin-sensitive-ATP] and [Oligomycin-sensitive-Fusion5], while these metrics linearly increase with each other within the grey cell population) (**Fig. 4C**, right). Bivariate analyses of [Oligomycin-sensitive-ATP] and [Oligomycin-sensitive-Fission] demonstrates that Oligomycin induces fission in all A2780-CPs, while Oligomycin can induce or suppress fission in A2780-IPs (p value obtained from Fisher’s exact test).

In summary, mito-SinCe^2^ analyses reveals that the mitochondrial dynamics status is dependent on mitochondrial energetics in the ovTIC^Aldh+^ enriched A2780-CPs in comparison to the parental A2780-IPs. It remains to be tested if and how mitochondrial dynamics modulates energetics in this system, which needs specific experimental design.

### Identification of cell populations with high or low mitochondrial potential (ΔΨ^hi/lo^) that equilibrate with each other in ovTICs^Aldh+^

Mito-SinCe^2^ analyses identified uniqueness in mitochondrial energetics in the ovTIC^Aldh+^ enriched A2780-CPs in comparison to the A2780-IPs (**Fig. 4**). Thus, we compared mitochondrial energetics in conventional ways between the A2780-IP and A2780-CPs in high density cultures that further enhances the expression of the ovTIC marker, Aldh1A (**Fig. S2D**). Firstly, bioenergetic profiling from mitochondrial OCR showed that the A2780-CPs have higher reserve respiratory capacity due to their elevated maximal respiration (**Fig. 5A,** details in Suppl methods). Secondly, flow-cytometric analyses of the A2780-CPs stained with potentiometric dye TMRE reveals a population with markedly low TMRE uptake, thus defining distinct ΔΨ^lo^ and ΔΨ^hi^ populations (**Fig. 5B**), which can be visibly confirmed in confocal micrographs (squares in **Fig. 5C**). The ΔΨ^lo^ peak is also detected in our newly derived paclitaxel-resistant A2780-PX line and a naturally CP resistant ovarian cancer line, SKOV-3 (Landen et al., 2010) (**Fig. S2E, F**). The non-potentiometric MTG stain does not detect distinct ΔΨ^lo^ and ΔΨ^hi^ populations (**Fig. S2E**) confirming that the ΔΨ^lo^ population is not due to reduced mitochondrial mass. Consistently, the dynamic range (obtained from single cells in confocal micrographs) of MTG or mitochondrial markers is 5-fold lower than that of TMRE that detects ΔΨ^lo^ cells (**Fig. S2G**).

**Fig. 5.**
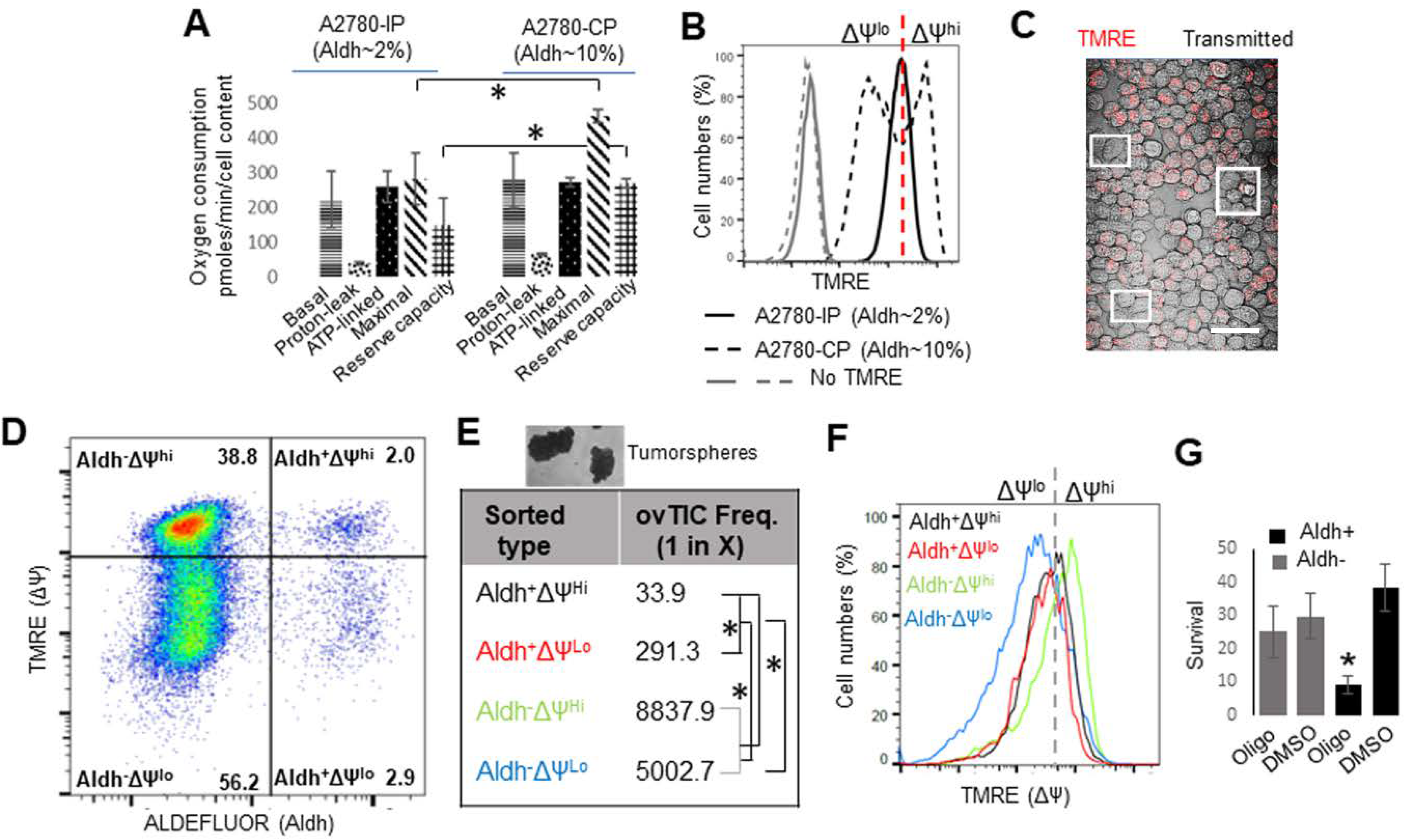
Identification of distinct ΔΨ^hi^ and ΔΨ^lo^ cell populations that equilibrate in ovTICs^Aldh+^. **(A)** Bioenergetic profiling of A2780-IP and A2780-CPs. * depict p <0.05 in T test. **(B)** Flow cytometric profile of TMRE stained A2780-IP and A2780-CPs. Dashed red line demarcates the ΔΨ^hi/lo^ population. **(C)** Confocal micrograph of TMRE stained A2780-CPs; boxes marking some ΔΨ^lo^ cells lacking TMRE and identified by transmitted light. Scale bar is 50 µm. **(D)** Representative bivariate plot of ALDEFLOUR and TMRE stained A2780-CPs; numbers reflect the percentage of cells. **(E)** Prediction of ovTIC frequency from ELDA of tumorsphere (image) forming ability of color coded cells sorted from A2780-CPs. **(F)** Flow cytometric profile of TMRE staining of color coded cells maintained in culture for 15 days in tumorsphere assay. Dashed grey line demarcates the ΔΨ^hi/lo^ cells. **(G)** Quantitation of survival of sorted Aldh^+^ and Aldh^−^ cells maintained in TIC enrichment conditions in the presence DMSO or Oligomycin. * depict p <0.05 in T test.

To determine if ΔΨ is linked to ovTIC self-renewal/proliferation, we took advantage of the previously reported TMRE based cell sorting (Schieke et al., 2008; Sukumar et al., 2016), and combined it to the ALDEFLUOR based ovTIC sorting. Using this approach, we isolated 4 populations, namely Aldh^+^ΔΨ^hi^, Aldh^+^ΔΨ^lo^, Aldh^−^ΔΨ^hi^ and Aldh^−^ΔΨ^lo^ (**Fig. 5D**). We used the standard extreme limiting dilution assay (ELDA) (Hu and Smyth, 2009) to determine ovTIC frequencies of each sorted population. Towards this end, we counted the frequency of tumorspheres formed in TIC medium and low attachment conditions that allow ovTICs to self-renew and proliferate *in vitro* (Schultz et al., 2016)(details of TIC medium in Methods). Expectedly, the ovTIC frequency is ∼200 fold higher in the Aldh^+^ populations than the Aldh^−^ populations, irrespective of ΔΨ (**Fig. 5E, Fig. S4H, I**). Interestingly, the ovTIC frequency in the Aldh^+^ΔΨ^hi^ population is ∼10 fold higher than the Aldh^+^ΔΨ^lo^ populations (**Fig. 5E, Fig. S2 H, I**). Importantly, the Aldh^+^ΔΨ^hi^ and Aldh^+^ΔΨ^lo^ populations equilibrate to form a population with an intermediate ΔΨ in the tumorspheres, while Aldh^−^ΔΨ^hi^ and Aldh^−^ΔΨ^lo^ maintain their original ΔΨ^hi^ or ΔΨ^lo^ status (**Fig. 5F);** the Aldh^+/−^ populations harbor proportionally comparable number of ΔΨ^hi/lo^ cells (**Fig. 5D**) and the Aldh^+^ status does not change during ΔΨ equilibration (not shown). Consistent with ΔΨ driving ATP synthesis, we investigated if the ovTICs^Aldh+^, which equilibrate between ΔΨ^lo^ and ΔΨ^hi^ states, are dependent on mitochondrial ATP synthesis as observed with certain other TICs (Viale et al., 2015). Indeed, 3-day exposure to picomolar dose of Oligomycin significantly reduced cell survival of Aldh^+^ cells, not affecting the Aldh^−^ cells (**Fig. 5G**).

Thus, we will employ mito-SinCe^2^ to investigate any existing linear relationships between mitochondrial dynamics and energetics in the ΔΨ^hi/lo^ populations that equilibrate in mitochondrial energy dependent ovTICs^Aldh+^ of the A2780-CPs.

### Quantification of the state of mitochondrial dynamics in single ΔΨ^hi/lo^ and Aldh^+/−^ cells

Here, we describe the differences in mitochondrial dynamics status in the identified ΔΨ^hi/lo^ and the sorted Aldh^+/−^ A2780-CPs, as detected by the validated [Fission], [Fusion5] and [Diameter] metrics; [Fusion5] is particularly optimal and thus chosen (**Fig. 1G, K**).

Since the ΔΨ^hi/lo^ cells can be distinguished in the confocal micrographs (**Fig. 5C**), we quantified the dynamics metrics using the MTG signal in TMRE and MTG co-stained cells (given the functional TMRE signal cannot be used for this purpose). We compared cells in glucose and galactose, since galactose elevates OCR (**Fig. S3A),** and stimulates mitochondrial fusion (**Fig. 1C, E**) (Mishra and Chan, 2016). Replacement of glucose with galactose reverts ΔΨ in both the ΔΨ^hi/lo^ groups within 2 hours (**Fig. 6A**), reducing the markedly high dynamic range of [ΔΨ] within the ΔΨ^lo^ group (**Fig. 6B**). Bivariate analyses of [Fission] and [Fusion] reveals the subset of ΔΨ^hi^ cells having higher [Fusion5] (and lower [Fission]) than the ΔΨ^lo^ group (**Fig. 6C,** left, arrow), contributing to overall increased [Fusion5] (and decreased [Fission]) (**Fig. S3C**). [Diameter] is modestly higher in the ΔΨ^hi^ group and is further increased by galactose (**Fig. 6C, Fig. S3C**). [Fission] decreases linearly with increase in [Fusion5] within both ΔΨ^hi/lo^ groups in the presence of glucose or galactose (**Fig. 6D,** left, ‘–’ signify inverse linear relationship). However, linear decrease in [Diameter] with increase in [Fusion5] is detected only within ΔΨ^lo^ group in galactose (**Fig. 6D,** right).

**Fig. 6.**
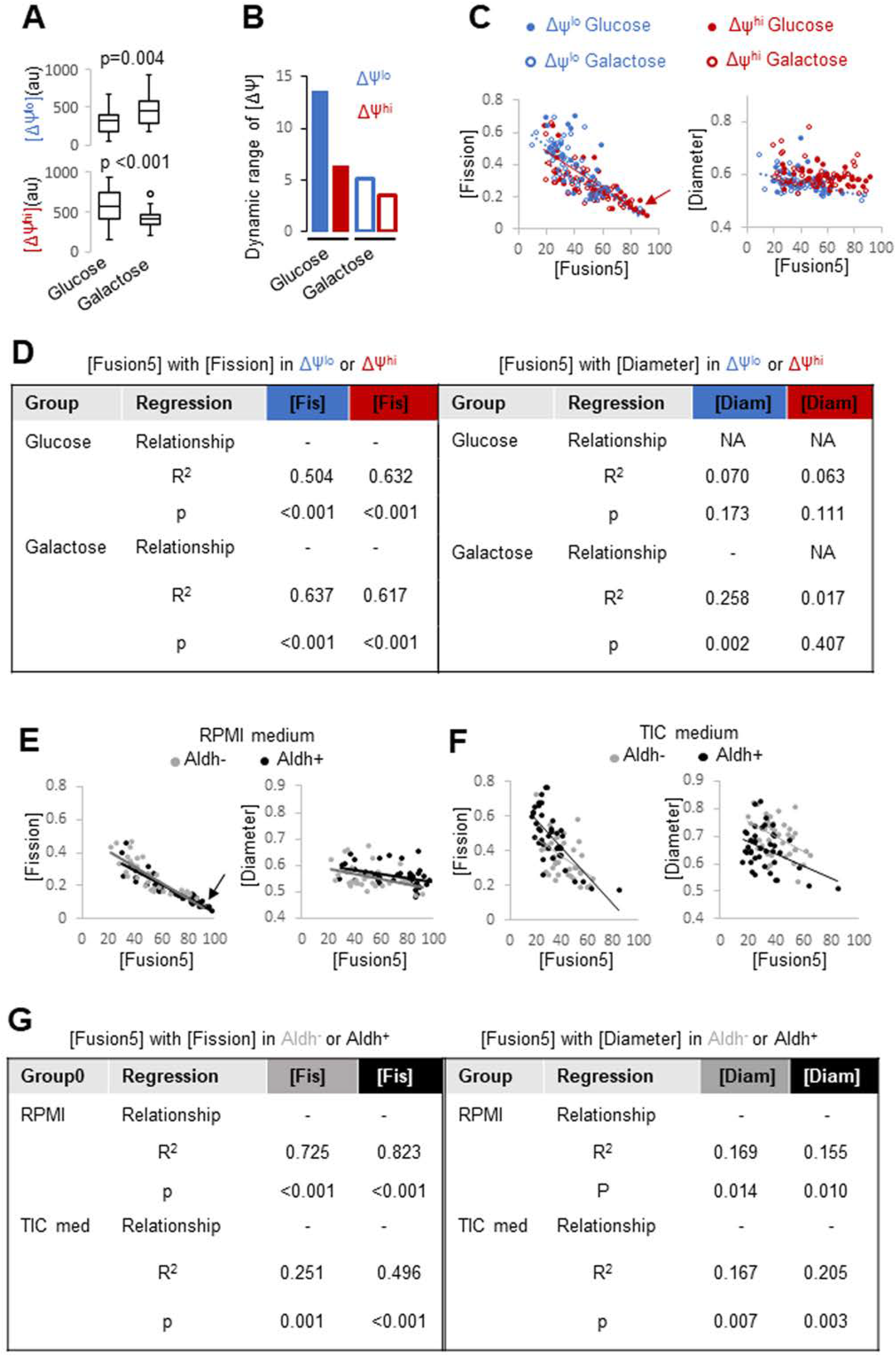
Quantification of the metric for dynamics in ΔΨ^hi/lo^ and Aldh^+/−^ populations. **(A)** Box-plots depicting quantification of TMRE (ΔΨ) signal from confocal micrographs of identified ΔΨ^hi^ and ΔΨ^lo^ groups in A2780-CPs maintained in glucose or galactose medium. The TMRE staining is not comparable between the ΔΨ^hi/lo^ group. p values are from non-parametric Kruskal-Wallis test. **(B)** Dynamic range of [ΔΨ] from (A). **(C)** Bivariate plot for regression analyses [Fission] or [Diameter] vs [Fusion5] in identified ΔΨ^hi/lo^ A2780-Cps maintained in glucose or galactose medium. Regression lines shown. Arrow points towards cells with >80 [Fusion5]. **(D)** R^2^, p values and relationship from regression analyses of metrics from **C**. NA: no statistically significant relationship. **(E)** Experiment in (C) performed in sorted Aldh^+^ and Aldh^−^ A2780-CPs, and thereafter maintained in RPMI medium. **(F)** Experiment in (E) in TIC medium. **(G)** Analyses as in (D) with data from (F).

Similar analyses of sorted and recovered Aldh^+/−^ cells stained with Mitotracker-633 (Suppl. Methods) revealed the subset of Aldh^+^ cells having higher [Fusion5] (and lower [Fission]) than the Aldh^−^ group in RPMI growth medium (**Fig. 6E**, arrow), contributing to the overall increase in [Fusion5] (and decrease in [Fission]) (**Fig. S3D**). However, TIC medium, designed to promote self-renewal and proliferation of ovTICs, maintains markedly higher [Fission] and lower [Fusion5] in comparison to the RPMI medium particularly in the Aldh^+^ group (**Fig. 6E, F, Fig. S3D**). TIC medium dramatically increased [Diameter] in both Aldh^+/−^ groups (**Fig. 6E,F, Fig. S3D**). [Fission] or [Diameter] increase linearly with decrease in [Fusion5] within both Aldh^+/−^ groups in RPMI and TIC media (**Fig. 6E-G**, ‘–’ signifies inverse linear relationship).

Therefore, ΔΨ^hi^ and Aldh^+^ groups harbor cells with unopposed fusion with >80 [Fusion5]. This implies that the Aldh^+^ΔΨ^hi^ group, which harbors maximum ovTIC frequency (**Fig. 5E**), has unopposed fusion state in RPMI medium that shifts to an unopposed fission state after exposure to TIC medium.

### Mito-SinCe^2^ analyses identifies the specific ΔΨ^lo^ population where the status of dynamics and ATP are linked

We employed the ATP arm of mito-SinCe^2^ (as in **Fig. 3**) to test if ΔΨ^lo/hi^ groups of the A2780-CPs have differential [Oligomycin-sensitive-ATP] in relation to the differential basal [Fission] or [Fusion5]. We identified ΔΨ^hi/lo^ groups in the TMRE stained mito-GO-ATeam2 expressing cells by generating [ΔΨ] histograms (Suppl. methods) (**Fig. S4A, B**).

K-means clustering analyses identified two distinct clusters in ΔΨ^lo^ group (predicted by gap statistics, see Suppl. Methods), which can be separated by [Fusion5] values into ΔΨ^lo^-Fusion5^hi^ and ΔΨ^lo^-Fusion5^lo^ clusters (**Fig. 7A**). ΔΨ^hi^ cells can be assigned to either of the two ΔΨ^lo^ clusters by their minimum Euclidean distance from the clustered ΔΨ^lo^ cells. As expected, similar assignment puts CCCP treated cells in the Fusion5^lo^ and the galactose utilizing cells in the Fusion5^hi^ clusters (**Fig. 7A**). Importantly, the ΔΨ^lo^–Fusion5^lo^ cells have markedly higher [Oligomycin-sensitive-ATP] in comparison to the ΔΨ^lo^-Fusion5^hi^ and ΔΨ^hi^ cells (**Fig. 7B**). Next, we performed linear regression analyses between parameters (**Fig. 7C, E**; ‘+’ or ‘–’ signify direct or inverse linear relationship, respectively; NA signifies no statistically significant linear relationship). Surprisingly, we detected no linear relationship between [Oligomycin-sensitive-ATP] and [Fusion5] within the ΔΨ^lo^–Fusion5^lo^ group that had maximum [Oligomycin-sensitive-ATP] (**Fig. 7C,** left). Only within the ΔΨ^lo^-Fusion^hi^ group, do [Oligomycin-sensitive-ATP] and [Fusion5] increase linearly with each other (**Fig. 7C,** right); [Fission] has opposite relationships than [Fusion5]. Notably, the [Fission] or [Fusion5] do not change linearly with ΔΨ within the ΔΨ^lo^ group (**Fig. S4C**). On the other hand, [ΔΨ] and [Fusion5] increase linearly with each other only within the ΔΨ^hi^ cells falling in the ΔΨ^lo^-Fusion^hi,^ cluster (**Fig. 7D**), and not within the larger ΔΨ^hi^ group (**Fig. S4D**).

**Fig. 7.**
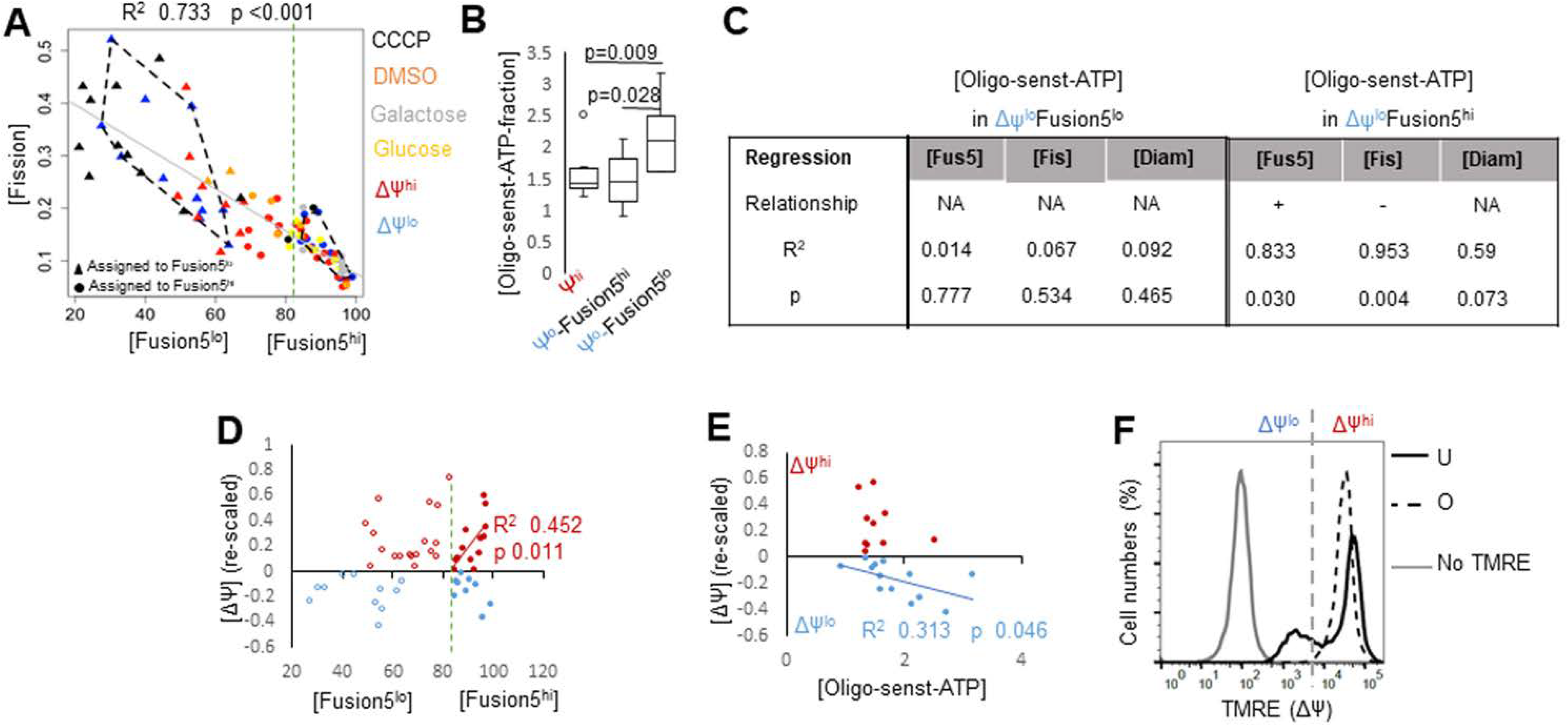
Mito-SinCe^2^ analyses of [ATP] vs. [dynamics] in single ΔΨ^hi/lo^ cells. **(A)** K-means cluster analyses of ΔΨ^hi/lo^ groups based on [Fission] and [Fusion5]; metric values of A2780-CPs maintained in Glucose, Galactose, DMSO or CCCP are overlapped on the identified clusters (demarcated with dashed lines); solid line represents the regression between [Fission] and [Fusion5] pooling all the groups. Dashed green lines demarcates Fusion-5^hi^ and Fusion-5^lo^ groups. **(B)** Box plots of [Oligomycin-sensitive-ATP] between the groups shown, while Fusion-5^hi/lo^ groups identified as in (A). p values are from non-parametric Kruskal-Wallis test. **(C)** R^2^, p values and relationship from regression analyses between shown mito-SinCe^2^ metrics in ΔΨ^lo^ group. NA: no statistically significant relationship. **(D)** Bivariate plot for regression analyses of [ΔΨ] and [Fusion5] in ΔΨ^hi/lo^ groups; dashed green line separates Fusion5^hi/lo^ groups. Only statistically significant regression line, R^2^ and p values shown. **(E)** Bivariate plot for regression analyses of [ΔΨ] and [Oligomycin-sensitive-ATP-fraction] in ΔΨ^hi/lo^ groups. Only statistically significant regression line, R^2^ and p values shown. **(F)** Flow cytometric profile of ΔΨ in TMRE stained A2780-CPs depicting the ΔΨ^hi/lo^ cell groups in untreated (U), treated with Oligomycin (O) or without TMRE; dashed gray line demarcates ΔΨ^hi/lo^ groups.

[Oligomycin-sensitive-ATP] is proportional to ATP synthesis that is proportional to ATP consumption, and modulated by ADP levels in steady state (Brand and Nicholls, 2011). Notably, enhancement of ATP synthesis reduces ΔΨ, provided the electron transport chain (ETC) activity is constant. Therefore, elevated [Oligomycin-sensitive-ATP] in the ΔΨ^lo^ group (**Fig. 7B**) can potentially reduce ΔΨ to maintain the ΔΨ^lo^ state. Indeed, ΔΨ linearly decreases with increase in [Oligomycin-sensitive-ATP] in the ΔΨ^lo^ group, but not in the ΔΨ^hi^ group (**Fig. 7E**). More importantly, inhibition of ATP synthesis by Oligomycin (10μM for 15 mins) increases [ΔΨ] of the ΔΨ^lo^ group, while slightly decreasing the ΔΨ^hi^ peak, which converts the ΔΨ^lo^-ΔΨ^hi^ groups to a single ΔΨ group (**Fig. 7F**). These results indicate that differential [Oligomycin-sensitive-ATP] may reflect differential ATP synthase activity between the ΔΨ^hi/lo^ group. Since Oligomycin also inhibits the reverse ATPase activity, the [Oligomycin-sensitive-ATP] reflects a smaller decrease than expected from ATP synthase inhibition.

In summary, quantitative mito-SinCe^2^ analyses reveals that a functional link between fusion and ATP synthesis is established beyond the value of [Fusion5]-80 in the ΔΨ^lo^-Fusion5^hi^ state. Such a level of fusion may represent a ‘hyperfused’ state that is linked to enhanced ATP production (Liesa and Shirihai, 2013; Schrepfer and Scorrano, 2016). Importantly, dynamics is not related to ATP in the ΔΨ^lo^-Fusion5^lo^ state although it maintains higher Oligomycin sensitive mitochondrial ATP. Also, the ΔΨ^hi^ state associates with an unopposed fusion state, while the ΔΨ^lo^ state may be maintained by elevated ATP synthesis (and minimum ETC regulation).

### Mito-SinCe^2^ analyses identifies the specific ΔΨ^hi/lo^ populations where the status of dynamics and redox are linked

We performed the redox arm of mito-SinCe^2^ (as in **Fig. 3**) to investigate if identified [ATP] vs. [dynamics] linear relationships (**Fig. 7C**) are linked to [redox] vs. [dynamics] linear relationships in the ΔΨ^lo/hi^ groups of A2780-CPs. Like in the HaCaTs (**Fig. 3E,** DMSO), regression analyses did not detect any linear relationship between [Oxidation] vs. [Fusion5 or Matrix-continuity] in the A2780-CPs (**Fig. 8A, F**). However, unlike HaCaTs, [Matrix-continuity] modestly decreases linearly with increase in [Fusion5] in the A2780-CPs (**Fig. 8B**) (this can arise due to discoordination between fusion of outer and inner mitochondrial membranes in the cancer line).

**Fig. 8.**
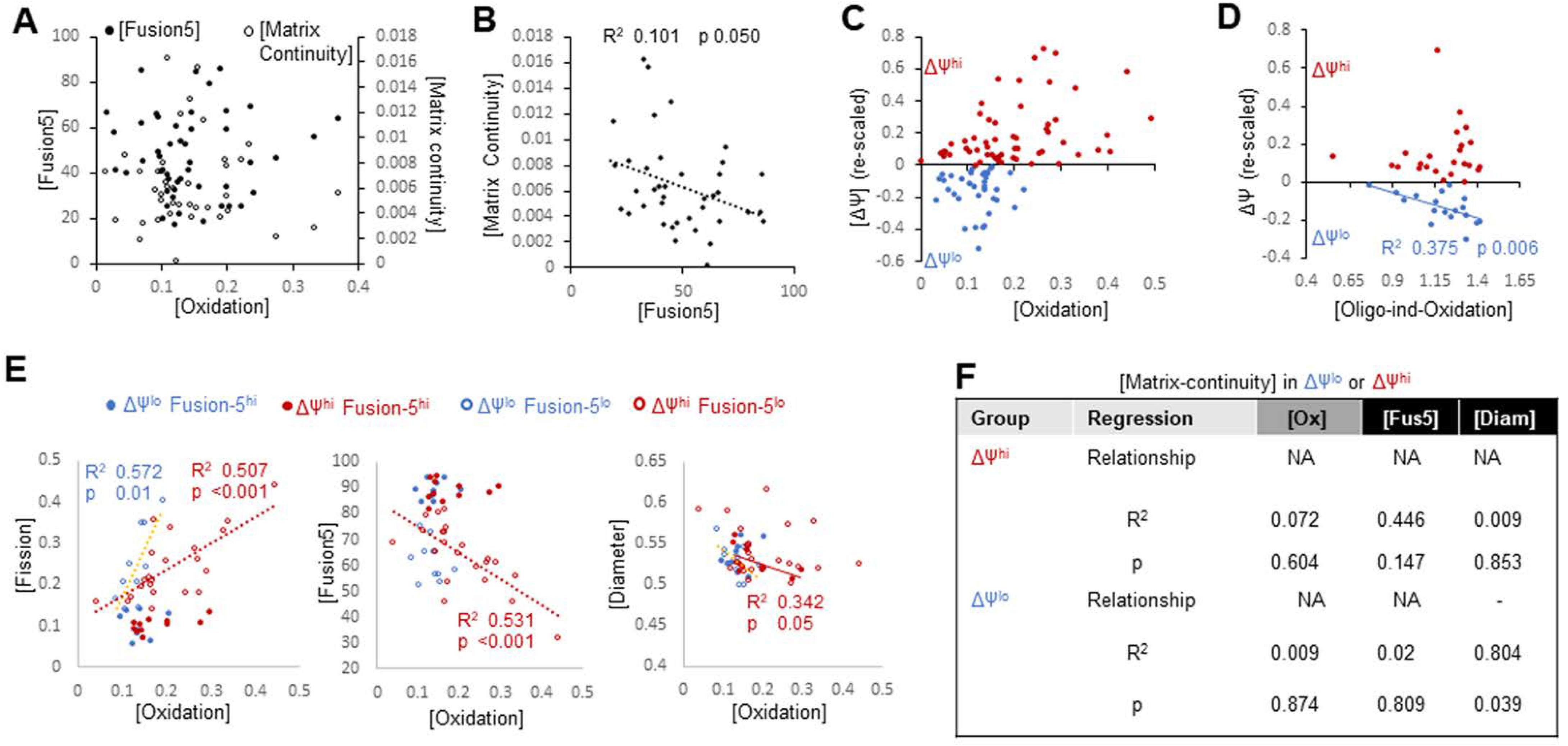
Mito-SinCe^2^ analyses of [redox] vs. [dynamics] in single ΔΨ^hi/lo^ cells. **(A)** Bivariate plot of [Oxidation] and [Fusion5 or Matrix-continuity] in A2780-CPs. **(B)** Bivariate plot of [Fusion5] and [Matrix-continuity] in A2780-CPs. R^2^, p value and regression line shown. **(C)** Bivariate plot of [Oxidation] and [ΔΨ] in ΔΨ^lo/hi^ groups. **(D)** Bivariate plot for regression analyses of [Oligomycin-sensitive-Oxidation] and [ΔΨ] in ΔΨ^lo/hi^ groups. R^2^, p value and regression line shown for only statistically significant regression. **(E)** Bivariate plot for regression analyses of [Oxidation] with dynamics metrics. Only statistically significant regression line, R^2^ and p values shown**. (F)** R^2^, p values and relationship from regression analyses between metrics shown in the ΔΨ^lo/hi^ group. NA.

**Fig. 9.**
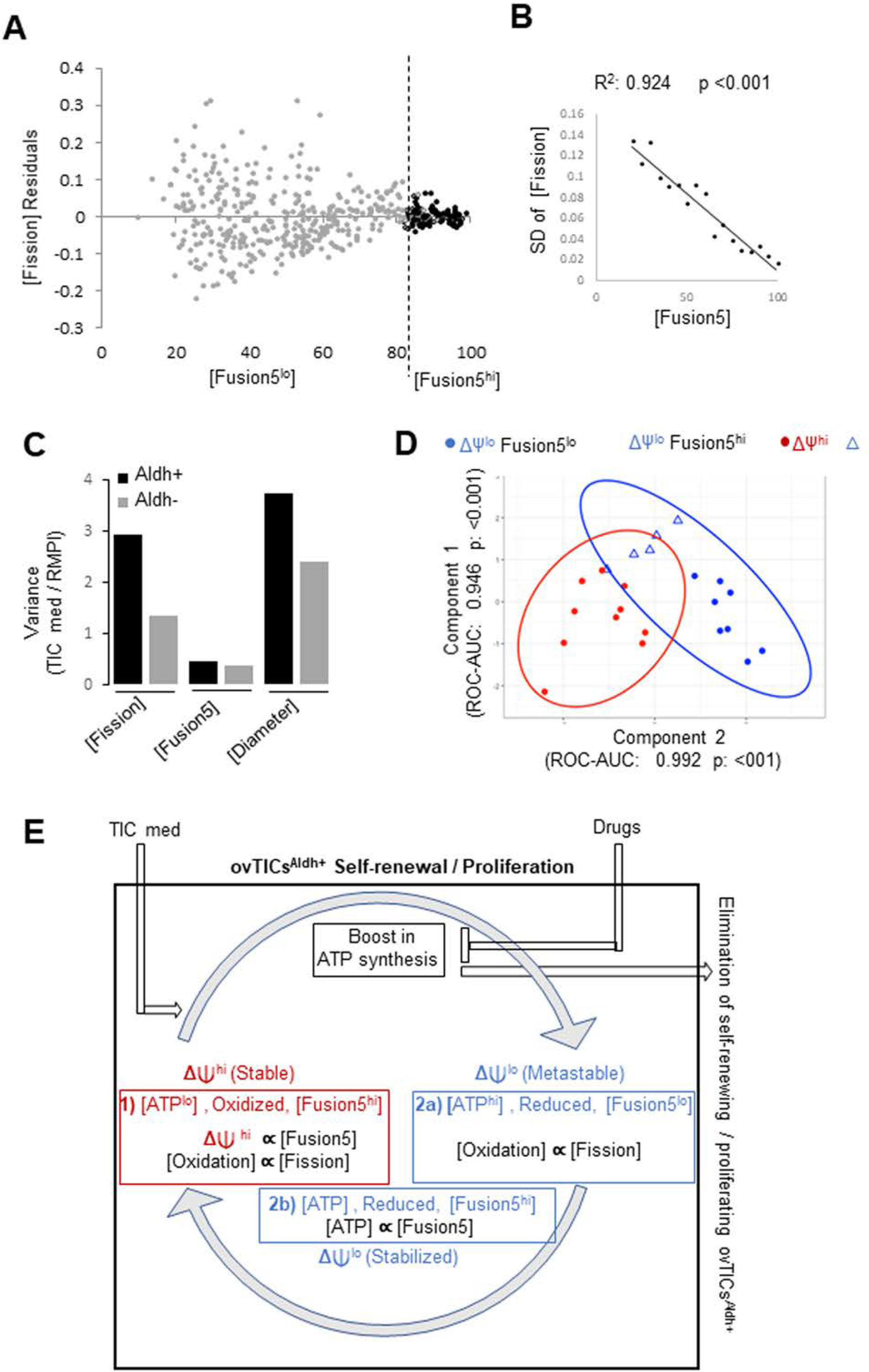
Potential predictive abilities of mito-SinCe^2^ about mitochondrial structure-function. **(A)** Bivariate plot of [Fission] residuals from various previous regression analyses with respective [Fusion5]; dashed line separates Fusion5^hi/lo^ cells in grey and black. **(B)** Regression of the standard deviation (SD) of [Fission] at progressively increasing windows of [Fusion5]. R^2^ and p values shown. **(C)** Comparison of the impact of variance of dynamics metrics between sorted Aldh^+^ and Aldh^−^ cells in RPMI or TIC medium. **(D)** Plot showing components 1 and 2 of PLSDA of the ΔΨ^lo/hi^ group. ROC-AUC and p values for clustering are shown. **(E)** Hypothetical model involving 3 distinct mito-SinCe^2^ states proposed for the underlying complexities of ovTIC^Aldh+^ energetics.

We identified the ΔΨ^hi/lo^ groups in mito-roGFP and mito-PSmO2 co-expressing cells modifying the strategy described in Fig. S4B **(Fig. S4E)**. Interestingly, [Oxidation] is maximum in ΔΨ^hi^ group (**Fig. 8C**), without a statistically significant linear relationship between [Oxidation] and ΔΨ in this group. Notably, in the ΔΨ^hi^ group the low [Oligomycin-sensitive-ATP] does not linearly change with [dynamics] **(Fig. S4D**). Although, Oligomycin (10 μM for 10 mins) oxidizes cells in both the ΔΨ^lo/hi^ groups (**Fig. S4F**), [Oligomycin-induced-Oxidation] linearly increases with decrease in [ΔΨ] only within ΔΨ^lo^ group (**Fig. 8D**), similar to increase in [Oligomycin-sensitive-ATP] (**Fig. 7F**). This raises the possibility that Oligomycin induced inhibition of ATP synthesis in the ΔΨ^lo^ cells dissipates the ΔΨ consumed for mitochondrial ATP synthesis to cause matrix oxidation.

Next, to analyze [Oxidation] vs. [dynamics] linear relationships, we partitioned the cells into Fusion5^hi/lo^, which differ in [Oligomycin-sensitive-ATP] and its relationship with dynamics (**Fig. 7A-C**). Importantly, [Oxidation] and [Fission] linearly increase with each other (**Fig. 6E**, left), within both ΔΨ^hi^Fusion^lo^ and ΔΨ^lo^Fusion^lo^ groups where no linear relationship was detected between [ATP] and [dynamics] (**Fig. 7C**). Notably, [Fusion5] linearly decreases with [Oxidation] only in the ΔΨ^hi^-Fusion^lo^ cells (**Fig. 8E,** middle). It remains to be seen, if enhanced reactive oxygen species induce fission in the Fusion^lo^ groups similar to what has been reported in some systems (Willems et al., 2015). We noted that [Diameter] linearly decreases with [Oxidation] within the ΔΨ^hi^-Fusion^hi^ group (**Fig. 8E,** right), where [ΔΨ] and [Fusion5] linearly increase with each other (**Fig. 5G**). This interesting connection between [ΔΨ], [Fusion5], [Diameter] and [Oxidation] in the ΔΨ^hi^-Fusion^hi^ group can be further explored in pathological situations associated with swelling of mitochondria. Such a conundrum does not exist in the in the ΔΨ^lo^ group where [Diameter] linearly increases with decrease in [Matrix-continuity] (**Fig. 8F**).

Thus, comparing data form **Fig. 7** and **Fig. 8**, we conclude that mitochondrial dynamics status changes linearly either with ATP or with redox, but not simultaneously with both. Also, the ΔΨ^hi^ cells have elevated diameter and maximally oxidized mitochondrial matrix in a fission dependent manner.

### Potential predictive abilities of mito-SinCe^2^

To test if mito-SinCe^2^ analyses can have any potential predicting abilities, we further analyzed the quantitative mito-SinCe^2^ metrics obtained from A2780-CPs described above.

[Fusion] and [Fission] may not always hold opposite relationship with energetics metrics (**Figs. 1L, 3E, 4C, D**). This highlights the distinction between the [Fusion] and [Fission] metrics and indicates they may not be interchangeably used although an inverse linear relationship exists between them. However, the inverse [Fission] vs. [Fusion5] linear relationship is not uniform between Fusion5^lo/hi^ cells, as reflected in the higher spread of data points from the regression line in the Fusion5^lo^ range (**Fig. 7A**); such non-uniformity in inverse [Fission] vs. [Fusion5] linear relationship was also observed in pooled data from various cell types (**Fig. S1L**). To probe further, we pooled data from **Figs. 6-8** to analyze the [Fission] residuals from linear regression analyses (indicating deviation from the regression line) separately in the Fusion5^lo^ and Fusion5^hi^ range. The [Fission] residuals progressively increase with decrease in [Fusion5] (**Fig. 9A**). with cells in CCCP and Galactose occupying zones with maximal and minimal residuals, respectively (**Fig. S4G**). Further analyses of standard deviation of [Fission] in windows of 5 [Fusion5] units revealed that the variation in [Fission] increases at a constant rate with decrease in [Fusion5] (**Fig. 9B**). Thus, this data predicts existence of factor(s) influencing the inverse [Fission] vs. [Fusion5] relationship.

TIC and RPMI media maintain distinctly different status of dynamics between the Aldh^+/−^ cells (**Fig. 6E-G**). TIC medium maintains higher variance in [Fission] specifically in the Aldh^+^ cells, while lower variance in [Fusion5] and higher variance in [Diameter] in both Aldh^+/−^ cells (**Fig. 9C**). This may predict that the TIC medium represses fusion in both Aldh^+/−^ cells, while maintaining a wider range of fission activity in the Aldh^+^ cells, similar to MFN1/2-DKO cells (**Fig. 1C**).

Next, we performed multivariate analyses to determine if combination of the mito-SinCe^2^ metrics can classify the cells in distinct ΔΨ^lo/hi^ clusters. Thus, we performed partial least square discriminant analyses (PLS-DA) using [Fission], [Fusion5], [Diameter], [Oligomycin-sensitive-ATP-fraction], and [ΔΨ]. The ΔΨ^lo/hi^ cells form statistically significant overlapping clusters with the majority of the cells in the non-overlapping zones and the overlapping zone consisting of the ΔΨ^lo^-Fusion^hi^ cells (**Fig. 9D**). While [ΔΨ] contributes maximally to the principal component 1 as expected, [Fusion5] contributes maximally to the principal component 2 (**Fig. S4H**). This confirms that the ΔΨ^lo/hi^ clusters are indeed distinct, while the overlapping zone may ‘potentially’ signify the path of equilibration of the two clusters as the ovTICs^Aldh+^ form tumorspheres (**Fig. 5F**). Thus, PLSDA of the mito-SinCe^2^ metrics of any unknown cell can be used to potentially classify the cell and predict its TIC and energetic state.

In summary, mito-SinCe^2^ can potentially make the following experimentally testable predictions: **a)** new features that can influence the fission-fusion relationship; **b)** if any physiological or pathological condition particularly impacts fission and/or fusion; **c)** classification of cells into functional classes.

## Discussion

The microscopy based high-resolution Mito-SinCe^2^ approach is the first step towards quantitative analyses of the interplay between mitochondrial dynamics (fission, fusion, matrix continuity and diameter) and energetics (ATP or redox) using single cells. We designed and validated individual mito-SinCe^2^ metrics (**Figs. 1,2**), provided proof of principal of the mito-SinCe^2^ approach by refining published findings (**Fig. 3**), employed the approach to address a biologically relevant question (**Fig. 4-8**), and generated predictions to be tested experimentally in future (**Fig. 9A-C**). The mito-SinCe^2^ approach can be used in specific experimental design to understand cause and effect relationship between dynamics and energetics (**Fig. 4**). Also, data generated from the mito-SinCe^2^ approach can be analyzed in various other statistical ways to refine the linear relationships reported here in or to identify meaningful non-linear relationships between the mito-SinCe^2^ metrics. Currently, mito-SinCe^2^ remains a low throughput approach and can be made high throughout with the development of high resolution automated confocal microscopes.

We employed mito-SinCe^2^ to investigate the distinct ΔΨ^lo^ and ΔΨ^hi^ cell groups that equilibrate in ovTICs^Aldh+^ (**Fig. 5F**) and likely maintain differential ATP synthesis (**Fig. 7B**). Based on mito-SinCe^2^ analyses we hypothesize that mitochondria-dependent self-renewing / proliferating ovTICs^Aldh+^ interconvert between 3 mito-SinCe^2^ states, namely State 1 (ΔΨ^hi^), State 2a (ΔΨ^lo−^Fusion5^lo^) and State 2b (ΔΨ^lo−^Fusion5^hi^) (**Fig. 9D)**. We speculate, that the reducing equivalents from carbon sources could build up ΔΨ in state 1, ATP synthesis in Fusion5^lo^ mitochondria with weakened energetic efficiency (Liesa and Shirihai, 2013) could dissipate ΔΨ in state 2a, and establishment of a direct relationship between ATP synthesis and fusion in state 2b could aid the transition from ΔΨ^lo^ to a ΔΨ^hi^ state. Importantly, dynamics relates to redox but not to ATP in states 1 and 2a, and to ATP but not to redox in state 2b.

Mito-SinCe^2^ analyses indicate how TIC self-renewal may involve both unopposed fusion and fission states that have been individually linked to stemness (Chen and Chan, 2017). We propose that during the cycling of the Aldh^+^ cells between ΔΨ^hi^ and ΔΨ^lo^ states (**Fig. 9E**), the Aldh^+^ΔΨ^hi^ cells with unopposed fusion state is ‘primed’ towards shifting to an unopposed fission state supported by the TIC medium (**Fig. 6E, F**). This explains the maximal ovTICs enrichment in the Aldh^+^ΔΨ^hi^ group (**Fig. 5E**); reduced ovTIC frequency of the Aldh^+^ΔΨ^lo^ group could be due to lag time for conversion to Aldh^+^ΔΨ^hi^ state through cell cycle transitions (not shown).

Regulation of TICs by mitochondrial energetics has been demonstrated in certain cases, but remains controversial (De Francesco et al., 2018). Our data indicates that interconversion between mitochondrial ATP^hi/lo^ states occurs during self-renewal and proliferation of ovTICs (**Fig. 9E**). Our data supports a concept where the ATP^hi^ΔΨ^lo^ state is metastable (likely due to elevated ATP synthesis with minimal ETC regulation), and the stabilization of this state is achieved by conversion to an ATP^lo^ΔΨ^hi^ state. The same phenomenon can be tested in other normal or neoplastic stem cells.

Currently, mitoSinCe^2^ can be used on immortalized/primary cells, including patient cells. Expression of relevant probes in transgenic animals will allow *in/ex vivo* mito-SinCe^2^ analyses. Appropriately targeted probe sets to other mitochondrial compartments would expand mito-SinCe^2^ abilities that currently focuses on the matrix. Finally, mito-SinCe^2^ analyses can be expanded to include other functional parameters like mitochondrial calcium, DNA, MAVs, moonlighting proteins on mitochondria etc.

## Methods (details in Suppl. Method)

### Cell lines, Reagents and Constructs

Cell lines were either purchased from ATCC or provided as gifts, while PX-resistant A2780 cell line was generated in house. Media and biochemical reagents were obtained from GIBCO or Sigma, ALDEFLOUR reagent from Stem Cell Technologies, Antibodies from BD Biosciences or Santa Cruz Biotechnologies. We sub-cloned the mito-roGFP (from Addgene) into pCDH lentiviral vector with hygromycin selection. We constructed the mito-PSmOrange in the lentiviral vector pCDH vector with puromycin selection, by tagging mitochondrial targeting sequence of the human cytochrome oxidase VIII subunit to the previously reported PSmO2 (gift from Dr. George Patterson). Mito-GO-Ateam2 is a gift from Dr. H. Imamura. Cell lines were maintained and transfections / transductions were carried out according to (Tanwar et al., 2016).

### Flow Cytometry and Cell Sorting

The ALDEFLUOR reagent was excited using a 488 nm laser with a 525/50 BP (505LP) filter on LSR II flow cytometer and a 530/30 BP (505 LP) filter on the FACS Aria II. TMRE was detected using a 532 nm laser and 582/15 BP filter on the LSR II and using a 561 nm laser and the 576/26 BP filter on the FACS Aria II. Cell sorting was performed after applying appropriate gating and compensations.

### Tumorsphere assay

Sorted cell groups were maintained in standard ovTIC medium that constitutes of RPMI (without TMRE stain) supplemented with N1 supplement, EGF, FGF, and Insulin. Limiting dilution spheroid assays were conducted by seeding 1, 10, 100, and 1000 cells in a 96-well ultra-low attachment plate with media supplemented every alternate day and counting spheres after 15 days. Then, the tumorspheres were dissociated by pipetting to analyze ALDEFLOUR/TMRE profile. Sphere formation was analyzed using the ELDA statistics.

### Microscopy

Cells were seeded in Labtek chambers (on Geltrex in case of sorted cells) at least 24 hrs before experimentation, and added with fresh medium 1 hr before microscopy.

Confocal microscopy of live cells was carried out on a laser scanning confocal microscope (LSM 700, Zeiss Microscopy) using a 40X Plan-apochromat 1.4NA/Oil objective, in a temperature and CO_2_ controlled chamber. Following lasers were used for excitation: 488 nm for mito-GO-ATeam2, 405nm and 488 nm for mito-roGFP, 555nm for basal, 488 nm for photo-switching and 639 nm for photo-switched mito-PSmO2. Appropriate detectors were used in each case. Multichannel image acquisition was designed to rule out crosstalk and cross excitations, with automated switching lasers between channels after each scanned line. 3D Confocal images were acquired at optical zoom 3 with 1 Airy unit pinhole and 0.5 µm Z interval.

For photo-switching mito-PSmO2 488 nm laser was used at 40-90% power and 2 iterations within a 50 by 50 px ROI. Thereafter, the photo-switched pool was followed by time lapse microscopy (2 µm optical slice) within 2 minutes at 15 second intervals.

Staining with TMRE (or with Mitotrackers) was carried out as previously described (Mitra and Lippincott-Schwartz, 2010). Higher concentration of TMRE and MTG and increased laser power were used for the ΔΨ^lo^ cells (for **Fig. 4**). To obtain the histogram profile of TMRE in cells expressing mito-GO-ATeam2 or mito-PSmO2 we designed a specific strategy, since TMRE spectrum overlaps with these probes. There, TMRE signal was re-scaled to convert the value of the inflection point of the ΔΨ^hi/lo^ peaks to ‘zero’.

To obtain the TMRE, ATP and redox status, maximum intensity projections of the optical slices were obtained, and appropriate ratios of the mean background corrected fluorescence were obtained from ROIs drawn around whole cells.

### MitoGraph v2.1 analysis

The MitoGraph v2.1 software involves an R script that uses the igraph package to calculate mitochondrial network properties per mito-component. MitoGraph v2.1 is freely available at https://github.com/vianamp/MitoGraph. MitoGraph v2.1 was run on TIFF images of individual cells with a single relevant fluorescent channel. The output files (.mitograph) were analyzed in Microsoft Excel using custom built macros.

### Immunoblotting and Colony Assays

These standard assays were carried out as previously described (Tanwar et al., 2016).

### Metabolic flux analyses

This was performed on Seahorse XF24 platform (Agilent). 25K (for **Fig.2**) or 100K cells were seeded and analyzed after 24 hours. After experimentation cells were immediately fixed by formalin and stained with crystal violet to get total cell content that was used to normalize OCR values. The bioenergetic profiling was performed as in (Brand and Nicholls, 2011).

## Statistical Analyses

Student’s t-test or Kruskal-Wallis tests, and linear regression analyses were performed with Microsoft Excel or SPSS package. K-means clustering, PLSDA and non-linear regression analyses, Fisher’s exact test, were performed using appropriate R packages. To rule out false positive discovery, our approach involved multiple testing correction for any regression analyses or box plot comparisons between two given parameters that were performed at least 10 times on any data set. Statistical significance was considered with p value <0.05.

## Author contributions

BS designed and performed experiments and analyzed data along with PG, AM, DP, MEF; ABH made constructs and helped in data analyses; MKB analyzed data; VDM provided consultation; KM conceived the study, helped in experimental design, analyzed data and wrote the manuscript with BS.

## Acknowledgements

BS, KM, DP are supported by NIH award (# R33ES025662). We acknowledge Drs G Patterson and H Imamura for sharing constructs, Drs K Mihara, D Chan and C Landen for sharing, Drs S. Rafelski and M. Vienna for help with MitoGraph version, J. Wirth for custom built excel macro, NIH grant supports P30 DK 079626 DRC for BARB Core and P30 AR048311 and P30 AI027667 for CFCC core.

## SUPPLEMENTAL INFORMATION

### 1. Supplemental methods

#### Cell lines

The A2780-IP and A2780-CP were gifted by Dr. C. Landen, The Drp-KO and its corresponding Wtd MEFs were gifted by Dr. K. Mihara and the MFN1/2-DKO MEFs were gifted by Dr. D. Chan.

#### Developing Paclitaxel-Resistant Lines

Paclitaxel resistant lines were developed by culturing parental A2780-IP ovarian cancer cells in 3 nM Paclitaxel. We seeded cells at roughly 40% confluence in 60mm dishes and added drug the next day. While cells were actively dying, we changed media regularly on every alternate day. After two weeks, resistant colonies started to emerge. We passaged cells into drug free media and added drug after attachment. We continue to maintain these cells in 3 nM Paclitaxel to prevent loss of resistance, but the cells are not exposed to the drug when they are seeded for an assay.

#### Flow Cytometry with AldeFluor

We used the AldeFluor reagent for activity of ovTIC marker aldehyde dehydrogenase (Aldh), which uses an Aldh substrate tagged to a fluorophore and the Aldh inhibitor DEAB to establish a baseline of fluorescence. We stained 2 million cells/mL with 5 µL AldeFluor reagent per mL AldeFluor Buffer. The inhibitor control was carried out by immediately transferring a portion of the stained cells into a tube containing DEAB to a final concentration of 10 µL inhibitor per 1 mL buffer. These were stained for 45 minutes at 37 C in 5% CO_2_. After staining, cells were centrifuged for 10 minutes at 300 g and re-suspended in AldeFluor buffer. When doing AldeFluor/TMRE assays, cells were re-suspended in room temperature buffer. Otherwise, cells were re-suspended in cold buffer and maintained on ice until completion of the assay. We confirmed that temperature did not impact AldeFluor fluorescence profile for our system provided the assay was carried out within 2 h (not shown).

A recovery period (∼12 hr) is necessary after Aldefluor sorting before performing assays to investigate mitochondrial properties because both FACS and Aldefluor staining impact mitochondrial morphology (data not shown).

#### Recognizing ΔΨ^hi/lo^ Populations on the Microscope

To obtain the histogram profile of mitochondrial membrane potential in cells expressing mito-GO-ATeam2 or mito-PSmO2 we designed the following strategy, since TMRE excitation and emission profile overlaps with these genetically encoded probes (**Fig. S4A,E**). Appropriate microscopy of the same cells was performed before and after TMRE staining. Hereafter, the fluorescence excited by 555 nm laser (FL_555_) of the pre-TMRE image of an individual cell was subtracted from the respective post-TMRE image to obtain the contribution from TMRE only. In this strategy, TMRE signal was saturated in the ΔΨ^hi^ group in the Mito-SinCe^2^ with mito-PSmO2 due to its higher laser power requirement, precluding any regression analyses in this group. After subtracting the mean EX-555 fluorescence intensity of the pre-image from that of the post-image, the values were plotted in a histogram with a bin-width of 200. We used the inflection point between the high and low peaks and the difference between the lowest and highest values to normalize these values with the inflection point set to zero. This allowed us to pool TMRE values from different days. The ratiometric analysis of the basal ATP and Redox probes were performed from the pre-TMRE images, to avoid any impact of TMRE on those mitochondrial functions. Because we found that Oligomycin ablates the ΔΨ^lo^ population, the Oligomycin sensitive ATP fraction had to be measured after TMRE staining. This precluded measuring Oligomycin driven changes in dynamics in these cells because TMRE, as a functional stain, reduces the accuracy of the MitoGraph v2.1 output.

#### MitoGraphv2.1 analyses

Individual cells were cropped and exported as stacked TIFFs with a single fluorescence channel. The images from a single experiment were saved to a folder on the Desktop. The MitoGraph software could then be run on these images by entering the following command string in Terminal:./MitoGraph –path ∼/Desktop/FolderName –xy 0.104 –z 0.5 –analyze. The MitoGraph files were pulled from the output and combined in Excel using a macro obtained from ExcelRibbon (https://excelribbon.tips.net/T010400_Importing_Multiple_Files_to_a_Single_Workbook.html) and modified by Jayson Wirth. After an initial quality control step (see below), the calculations of [Fission], [Fusion(1-10)], and [Diameter] were performed in Excel. Elements not assigned volumes were disregarded for purposes of [Diameter] though not for [Fission]. If the sum of these percentages was more than 0.005 off of 100% of the total volume reported by MitoGraph, this volume was replaced by the sum of the volumes reported for each individual element.

For quality control the binary MitoGraph outputs were compared to the TIFFs to determine whether coverage was acceptable in randomly chosen cells. Binary images were definitively checked if the MitoGraph output said the cell: 1) had a total length of less than 100 um, 2) had a total component number of less than 30 or greater than 100, or 3) had a longest fragment of less than 20 µm. After acquiring our metrics, the binary images of outliers were reexamined. In the analyses using the Mitotrackers with TMRE (**Fig. S3B**), cells with an [Fission] of greater than 0.7 were excluded because of concerns about the health of these cells, but their inclusion does not alter our conclusions.

#### K-means clustering

K-means clustering was carried out using the “cluster” package in R using default parameters. The number of clusters (*k*) was determined using *clusGap* function of the R package “*cluster*” and with visual examination using *fviz_nbclust* function from *factoextra* package. Briefly, the “gap statistics” of a cluster is the change of variance of a random set points within the cluster boundaries and variance of the actual points, after clustering the points into k clusters. This statistic is calculated for different values of k. The final number of cluster was determined using the the default method of “firstSEMax”. In this method, the optimum number of k is the smallest value of k for which the gap statistic value is within 1 standard error of the lowest local maximum value.

The other points were assigned a cluster based on the average euclidean distance of each point from the points in each magenta cluster. A point was assigned a cluster that has the smallest average distance from this particular point to all the members of the cluster.

#### PLS-DA

Partial Least Square Discriminant Analysis (PLS-DA) is used to classify points based using partial least square regression. The analyses were carried out using R *MixOmics* package.

#### Non-linear curve fitting

We used Generalized Additive Model (GAM; 1,2) to identify the best-fit curve for data showing non-linear relationship. We fit cubic spline to the dataset using the “*gam*” function from the R package *mgcv* (3) for this purpose. The R^2^ estimate of each model was determined using the proportion of the variance explained by the fit as implemented in mgcv package.

#### Outlier detection

For detection of outliers in a dataset, we used the standard interquartile range (IQR). In each dataset we subtracted the lower quartile from the upper quartile to calculate the IQR. Points above upper quartile or below lower quartile beyond 1.5 times of IQR was considered outliers.

#### Photo-switching based assay for matrix continuity

We thoroughly characterized our newly constructed mito-PSmO2 probe. HEK cells expressing mito-PSmO2, when stained with MTG, exhibited co-localization of mito-PSmO2 with MTG. This confirmed the anticipated localization of the mito-PSmO2 probe in the mitochondrial matrix (not shown). PSmO2 has a basal 555 nm laser excitable fluorescence that can be photo-converted with 488 nm laser to fluorescence excitable with 633 nm laser. Optimal photo-conversion (∼10 fold) of the basal fluorescence to the activated fluorescence was achieved by 90% 488 nm laser power with 2 iterations on a laser scanning confocal microscope. Only 3 fold or more photo-converted pools were considered for quantification. Cells expressing mito-PSmO2 can be locally photo-converted to highlight a mitochondrial pool (pulse). The highlighted mitochondrial pool in the region of interest gets diluted overtime and the basal pool recovers within 120 secs primarily due to matrix continuity. The ratio of the increase in basal fluorescence and concomitant decrease in photo-converted fluorescence is quantified, with appropriate background and bleaching corrections, by determining the linear coefficient of the parabolic fit to the time lapse.

Photo-activatable probes have been previously used to semi-quantitatively assess fusion by monitoring the dilution of the photo-activated pool due to active fusion over 30 minutes (Karbowski et al., 2004) (Mishra et al., 2014; Xie et al., 2015). Our assay for quantification of [Matrix-continuity] is distinct from these reported assays of fusion in that it measures recovery of fluorescence over 2 mins and controls for physical movement of mitochondria in and out of the regions of interest (ROI), indicated by decrease in both basal and photo-switched pools **(Fig. S1C,** dashed lines).

It is noteworthy that the photo-converting 488 laser also bleaches mito-roGFP signal. Thus, our analyses of Redox states is consistently from pre photo-converted images. Neither mito-PsmO2 nor mito-roGFP can be used with mito-GO-ATeam2 due to overlapping fluorescence spectra so the matrix continuity assay is only compatible with the Redox arm of Mito-SinCe^2^

#### Composition and use of TIC media

For TIC media, we used a recipe (Schultz et al., 2016) designed to maintain proliferation and self-renewal of ovTICs while killing or inhibiting proliferation of differentiated tumor cells. To do this we added growth factors to a serum-free RPMI base. Specifically, we added 1x N1 supplement, 500 mg/mL insulin, 20 ng/mL EGF, and 10 ng/mL bFGF to a volume of 200 or 500 mL RPMI with 1% sodium pyruvate, 1% L-glutamine, and 1% pen-strep. Aliquots of the growth factors were stored at −20C. We discarded unused media after 1 month and made fresh.

For the tumorsphere assay, we sorted cells into this TIC media, pelleted, and resuspended at the dilution required for the ELDA assay. We seeded the sorted cells on low attachment plates and every alternate day supplemented with TIC media equaling 10% of the volume in the wells.

For microscopy and survival/proliferation assays we handled sorted cells as above and then allowed them to recover overnight on Geltrex in the presence of growth medium (RPMI) or TIC medium because Geltrex maintains Aldh activity similar to suspension (data not shown). For experiments spanning several days, we replaced the total volume of the TIC media every alternate day. For microscopy experiments we seeded cells on glass bottom live cell chambers and replaced media at least 1 hr before beginning imaging. For survival/proliferation assays, we seeded cells densely into a six-well plate after sorting. When cells reached confluency, we replaced the media in each well with the appropriate media (TIC or growth) containing Oligomycin (250 pM) or DMSO (250 pM) and after 3 days, we fixed cells in formalin and stained with Crystal Violet for colorimetric quantification.

#### Bioenergetic profiling using Seahorse metabolic flux assay (Brand and Nicholls, 2011)

Non-mitochondrial Respiration: OCR independent of ETC activity, for instance by NADPH Oxidases. Obtained by measuring OCR after inhibition of ETC, e.g. inhibition of Complex III by Antimycin, and then adjusting by cell content.

Basal Respiration: Mitochondrial OCR under normal culture conditions. Primarily related to ATP production but also substrate oxidation and proton leak. Obtained by subtracting Non-mitochondrial Respiration from OCR in normal conditions (adjusted for cell content).

Proton Leak: Mitochondrial OCR that is not related to ATP-production. Related to production of reactive oxygen species by ETC activity. Obtained by adjusting OCR values by cell content and subtracting Non-mitochondrial Respiration from OCR after inhibition of ATP synthase, e.g. with Oligomycin (adjusted for cell content).

ATP-Linked Respiration: Mitochondrial OCR related to ATP-production by the mitochondrial ATP synthase. Obtained by subtracting Proton Leak from Basal Respiration. We found that ATP-Linked Respiration is positively correlated with the mean [CCCP sensitive ATP] in the cell lines we investigated (**Fig. 3C**). Here, we reported ATP-linked OCR as a percent reduction rather than as absolute values.

Maximal Respiration: The maximum OCR mitochondria in particular cells can achieve, for instance when under acute stress. Obtained by subtracting Non-mitochondrial Respiration from the OCR achieved on uncoupling the ETS from ATP synthase, e.g. using CCCP (adjusted for cell content).

Reserve Capacity: A measure of the extra work mitochondria could perform under acute stress conditions. The difference between the Maximal Respiration and the Basal Respiration. We found that when seeded densely the drug resistant A2780-CP line had significantly higher Reserve Capacity than the parental A2780-IP line (**Fig. 5A**).

We additionally computed the fold-change in OCR in response to inhibitors (Oligomycin, CCCP, and Antimycin) in order to report values analogous to those obtained from Mito-SinCe^2^ (**Fig. S1K**).

### 2. Supplemental results

#### MitoGraph v2.1 analyses of Cells 1-4

In addition to the functionalities available in v2.0, the MitoGraph V 2.1 yields the length and volume of individual mitochondrial components and total component number from high resolution 3-D confocal micrographs of live single cells. To test if the quantitative output from the MitoGraph v2.1 corroborated observed qualitative differences we selected four representative cells 1-4 (stained with Mitotracker green) with visually different mitochondrial morphology a cells (**Fig. 1A**). We found: **a)** cells 1and 2 have less than 50 whereas cells 3 and 4 have greater than 100 individual mitochondrial components (**Fig. S1A, X axis**); **b)** Cells 1 and 2 have their longest element constituting greater than, and Cell 3 and 4 less than, 20% of the total mitochondrial length (**Fig. S1A, Y axis**); **c)** Cell 1 and 3 harbor both long and short mitochondria, while some mitochondria have diameter greater than value 1 au (we refrain from using the unit as ‘µm’ since the derivation of diameter values from the reported volume involves certain assumptions) (**Fig. S1B**).

We derived the [Diameter] metric by weighting each element by its contribution to the total mitochondrial volume, analogous to the calculation of atomic weight by isotope abundance. We calculated the diameter of each element (d_n_) from its volume and length by assuming the elements are cylindrical, which should hold for most situations. We weighted each element with its percentage of the total mitochondrial volume (V_n_ / V_T_) and calculated the weighted mean diameter ([Diameter]) of the mitochondria for each cell 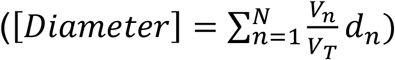. The weighted mean diameter, unlike the width output of MitoGraph v2.1, matched our subjective analyses and we proceeded with this metric.

#### Cautionary note emerging from our analyses of TMRE/MTG stained cells

We noted that glucose utilizing ΔΨ^hi/lo^ populations, stained with both TMRE and MTG, harbored a population of cells with [Fission] greater than 0.7 that had unexpected or no relationship with the [Fusion(1,5)] metric (**Fig. S3B**); these cells also had markedly higher average diameter (not shown). Cells with such high [Fission] are extremely rare as detected with MTG alone (even in MFN1/2-DKO MEFs), and their numbers were reduced by replacing glucose with galactose that stimulates mitochondrial oxygen consumption. Thus, we categorized the cells with >0.7 [Fission] as unhealthy, and filtered them out of the data set; this compromise in health may have resulted due to dual staining with TMRE/MTG. Nonetheless, including these cells in our analyses does not alter our overall conclusions.

#### Basic characterization of mito-GO-ATeam2 Specks

We noted that the mito-GO-ATeam2 ATP probe localization is not uniform in 10% of the overall cell population, with ∼5% in ΔΨ^hi^ and ∼8 % in ΔΨ^lo^ population. The fluorescence of the probe is enriched in specific regions that co-localize with Mitotracker 633 (data not shown) by significant recovery of fluorescence signal within 30 secs of bleaching in a Fluorescence Recovery After Photobleaching assay (Mitra and Lippincott-Schwartz, 2010) (**Fig. S5A**, arrow showing bleaching area). Although the mito-GO-ATeam2 specks reorganize in space and time similar to the rest of the mitochondrial population, they do not readily assemble or disassemble (data not shown). Analyzing a stable mito-GO-ATeam2 speck that did not change shape during the experimental time period, we found that although the [basal ATP] appears to be marginally higher in the mito-GO-ATeam2 specks, the [Oligomycin sensitive ATP] is lower than the uniform non-speck region in the vicinity (example in **Fig. S5B**). Additionally, when we include cells with these specks in our energetics analyses, we see markedly lower [Basal-ATP] and [Inhibitor-sensitive-ATP-fraction] in the MFN1/2-DKO MEFs than in WTm (data not shown), which we do not see in the filtered results. This suggested that the mito-GO-ATeam2 specks are bio-energetically less active with respect to mitochondrial ATP production. Consistently, exposure to the carbon source galactose reduces the number of cells with mito-GO-ATeam2 specks by 50% (**Fig. S5C**); inhibition of ATP synthesis by Oligomycin does not affect the abundance of the mito-GO-ATeam2 specks. We have also detected intracellular mitochondrial heterogeneity with the mito-roGFP probe, but in lower frequencies (not shown).

We excluded cells containing these specks from our analyses as they need further characterization. Additionally, they interfere with MitoGraph-based quantification (see minor limitations below).

### 3. Supplemental discussion

#### Minor limitations of MitoGraph-derived metrics

##### Mito-SinCe^2^ does not address the kinetics of fission and fusion

The mito-SinCe^2^ method and our [dynamics] metrics were designed to investigate the structure and function relationships of mitochondria across a range of cells and conditions, but we do not quantify active fission or fusion, only the fission or fusion state at given time points. These states reflect the fission and fusion activity but do not quantify kinetics and do not parse the regulation of fission and fusion by e.g. localization mechanisms from regulation for altering energetics. Additionally, because [Fusion(1-10)] are derived from mitochondrial length, they do not reflect transient fusion events that do not result in change in mitochondrial shape.

##### Impact of mitochondrial turnover

Our metrics attempt to account for mitochondrial turnover by controlling for total mitochondrial content, but the specific mechanisms of mitophagy and mitochondrial biogenesis can still influence our [Fission] and [Fusion(1-10)] parameters. Particularly where we see a breakdown of the fission/fusion relationship with our metrics, this could be related to turnover rather than fission and fusion, *per se*.

##### Impact of image resolution

MitoGraph v2.1, as with all image analysis software, is limited by the resolution of the image. Our resolution here is such that [Fission] may under-report fission (and [Fusion(1-10)] over-report fusion) in cells where mitochondrial components are maintained close together in space (e.g. perinuclear) but are still highly fragmented. Additionally, our resolution in the z-dimension may lead to a universal overestimation of average diameter, while still allowing us to compare between cells and treatments. For this reason, we refrain from reporting [Diameter] as µm, opting instead for arbitrary units (au) to make clear we do not expect [Diameter] to accurately report the width of mitochondria in µm but only to reflect differences in mitochondrial width among conditions and cell populations.

##### Overestimation of [Diameter] of small components

In MitoGraph v2.1, some of the smallest mitochondrial components are reduced to a single pixel by skeletonizing. To preclude these being treated as a node with no edges, they are arbitrarily given at least one neighbor on the xy plane. In cases where these components are elongated along the xz plane, the edge that was imposed on xy may not represent the major axis of the small mitochondrial component, meaning that for the smallest components, particularly in rounded cells, the [Diameter] might be overestimating the comparative average thickness. This will usually be minor because the average is weighted by volume, but may be important where small mitochondria make up a large fraction of the total mass. Where the impact is not negligible, these components are removed from the [Diameter] calculation.

##### Fluorescence intensity of dynamics probes must be similar across mitochondria within a cell

The relationship between [Fission] and [Fusion(1-10)] obtained from MitoGraph v2.1 output is optimal when there is roughly even fluorescence intensity. This is why functional probes which may vary in intensity within a cell if there is mitochondrial heterogeneity yield less accurate values. Additionally, mitochondrial structures which may impact mitochondrial morphology but which cause variation in fluorescence intensity of probes may interfere with MitoGraph v2.1. We have seen this in the bright specks in some cells expressing mito-GO-ATeam 2 (described above) as well as in the Drp1 knock-out MEFs, which show swollen regions within the mitochondrial network (**Fig. 2E**) that fluoresce so intensely that MitoGraph v2.1 misses thin linking regions, causing [Fusion(1-10)] to underestimate fusion in these cells.

##### Impact of particular probes

Comparison of the dynamics metric values obtained by using Mitotracker-633 and mito-GO-ATeam2 on Wtd / Drp1-KI and Wtm / MEN1/2-DKO MEF pairs reveal that the metric values may differ with probes, although the results are by and large consistent between the probes; mito-GO-ATeam2 appears to be more sensitive than Mitotracker-633. Particular fluorophores may impact mitochondrial morphology in ways that alter our metrics. Our quantitative analyses indicated a possible alteration of dynamics in cells doubly stained with TMRE/MTG (details in Suppl. Results, **Fig. S3B**), causing us to exclude certain cells from our analyses. Additionally, we have seen higher values for [Fusion(1-10)] in cells stably expressing mito-roGFP and higher [Diameter] in MFN1/2-DKO MEFs stained with MT 633 than those expressing genetically encoded fluorophores (this is not the case in the Drp1-KO MEFs). The possible impact of the particular probe on mitochondrial morphology must be considered when drawing conclusions from mito-SinCe^2^.

##### Possible impact of different MitoGraph versions

These analyses were performed with MitoGraph v2.1. A new version, MitoGraph v3.0 is now available. The primary differences between these versions of the software relate to *post facto* utility, but use of MitoGraph v3.0 will allow additional parameters to be considered.

## 5. Supplemental figure legends

**Figure S1.**
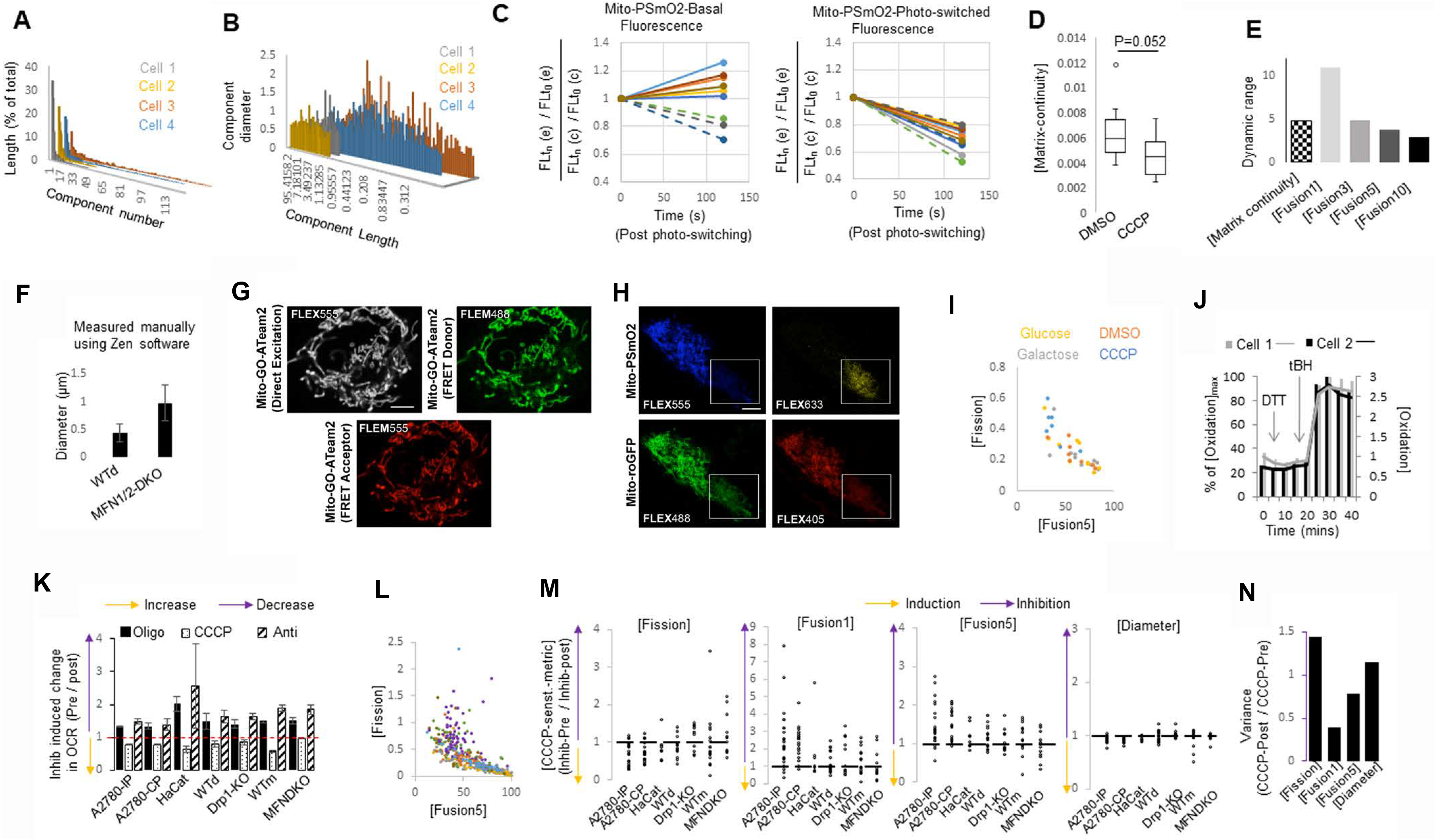
(A) Bar plot of length of individual mitochondrial component obtained using MitoGraph v2.1 on confocal micrographs of color coded cells (1-4) presented in main **Fig.1A**. The mitochondrial length of individual components is expressed as percentage of total mitochondrial length, and plotted in a descending order of length. Text describing results in Supplementary results section. **(B)** Bar plot of diameter of individual mitochondrial component derived from the volume obtained using MitoGraph v2.1 on confocal micrographs of color coded cells (1-4) presented in main **Fig.1A**. The diameter is plotted in a descending order of length of the individual mitochondrial component. Text describing results in Supplementary results section. **(C)** Lines plots depicting quantification of fluorescence signal from a photo-conversion based pulse chase assay with mito-PSmO2, where a photo-switched pool (pulse) is followed over time (chase). Normalized fluorescence of the basal pool (left panel) and the photo-switched pool (right panel) are shown. Color coded lines represent individual cells. Dashed lines represent the cells where both basal and photo-switched signal decreased with time, representing artifacts and thus excluded from further analyses. FL: Fluorescence intensity, tn: time point n, e: experimental cell, c: control cell for bleaching. **(D)** Box plot showing [Matrix-continuity] in HaCaT cells in DMSO or CCCP. p values are from Kruskal-Wallis test. **(E)** Bar plot showing dynamic range of the single cell metrics for dynamics obtained in Fig. 1K,L. **(F)** Bar plot showing the mean diameters of WT and MFN1/2-DKO MEFs, measured manually using the image acquisition software, Zen. **(G)** Representative confocal micrographs of the A2780-CP cells expressing mito-GO-ATeam2 FRET probe. FL_488_ channel is emission from the FRET donor excited with 488 nm laser, FL_555_ channel is from the FRET acceptor excited with 488 nm laser, FL_555Dir_ is from direct excitation of the FRET acceptor with the 555 nm laser. Scale bar is 5 µm. **(H)** Representative confocal micrographs of the A2780-CP cells expressing mito-roGFP and mito-PSmO2 probe. Micrograph represents image after photo-switching of the mito-PSmO2 that bleaches also mito-roGFP signal (our analyses of redox states from mito-roGFP is consistently from pre photo-converted images). FL_405_ channel is emission from excitation of mito-roGFP with 488 nm laser, FL_488_ channel is from excitation with mito-roGFP with 405 laser, FL_555_ is from excitation of mito-PSmO2 with 555 laser, FL_633_ is from excitation of the photo-converted pool of mito-PSmO2 with 633 nm laser. Scale bar is 5 µm. **(I)** Bivariate plot of [Fission] and [Fusion5] in HaCaT cells maintained in fresh Glucose, Galactose, DMSO or CCCP for 2 hrs. Metric values were obtained using FL_488_ of the mito-roGFP probe in the MitoGraph v2.1 analyses. **(J)** Bar plot showing impact of reducing agent DTT and oxidizing agent t-BH on Cell 1 and Cell 2 expressing mito-roGFP. [Oxidation] values represent raw data obtained from mito-roGFP fluorescence, while % of [Oxidation]_max_ represents percentage of maximal oxidation induced by t-BH. **(K)** Bar plot showing Oligomycin, CCCP or Antimycin induced change in OCR in 7 cell lines tested. **(L)** Bivariate plot of [Fission] and [Fusion5] in 7 cell lines tested. **(M)** Scatter plots showing quantitative changes induced by CCCP on the dynamics metrics in single cells in 7 cell lines tested. Dashed line represents no change. **(N)** Bar plot showing variance of the dynamics metrics from G. Variance calculated from data pooled from 7 cell lines tested.

**Figure S2.**
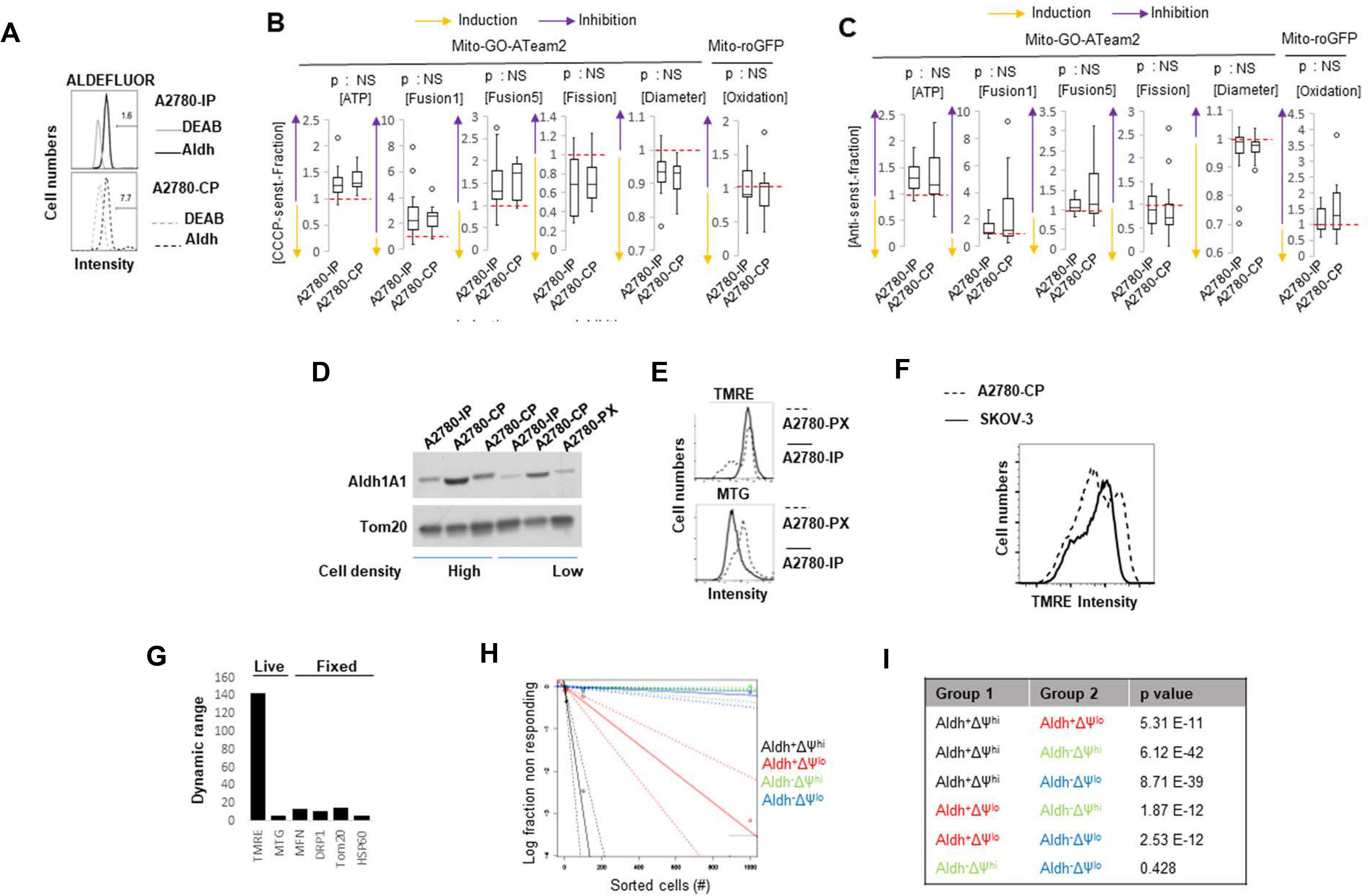
(A) Flow cytometric histogram profile of ALDEFLOUR stained A2780-IP and A2780-CP cells. DEAB treated cells serve as controls since DEAB inhibits Aldh activity. Numbers represent percentage of cells in the marked area of the histogram profile. **(B)** Box plots quantifying CCCP induced changes of energetics and dynamics in A2780 cells. Dashed lines represent no change. **(C)** Box plots quantifying Antimycin induced changes of energetics and dynamics in A2780 cells. Dashed lines represent no change. **(D)** Immunoblot analysis of Aldh1A1 in A2780-IP, A2780-CP and A2780-PX cells in high and low cell densities. Tom20 serves as a loading control. **(E)** Flow cytometric histogram profile of TMRE (upper panel) or MTG (lower panel) stained parental and derived A2780-PX cells. **(F)** Comparison of flow cytometric histogram profile of TMRE stained derived A2780-CP cells and naturally Cp-resistant SKOV-3 cells. **(G)** Bar plot showing dynamic range of TMRE, MTG, Drp1, Mfn1, Tom20 and HSP-60 staining in the A2780-CP cells. **(H)** ELDA plot of tumorsphere forming ability towards predicting the ovTIC frequency of the color coded sorted population of the derived A2780-CP cells. The ovTIC frequencies obtained are presented in main **Fig. 5E**. **(I)** Table showing the p values obtained from pairwise (Group 1 vs. Group2) Chi-Square test in ELDA analyses from (E).

**Figure S3.**
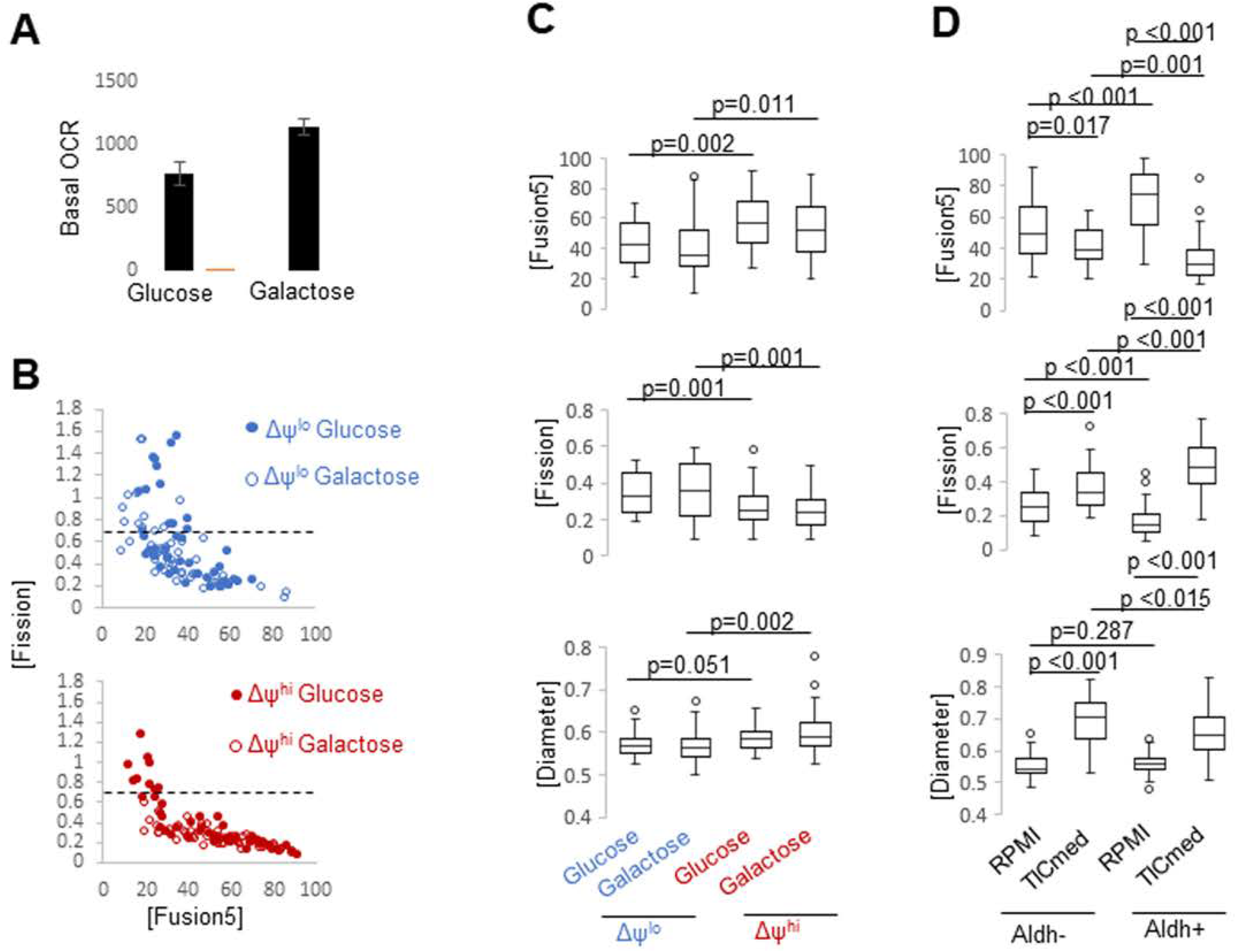
(A) Bar plot showing mean OCR in A2780-CP cells maintained in fresh Glucose or Galactose medium for 2 hrs. **(B)** Bivariate plot of [Fission] and [Fusion5] in TMRE and MTG stained A2780-CP cells, identified as ΔΨ^lo^ (upper panel) or ΔΨ^hi^ (lower panel) based on distinct differences in their TMRE uptake. Cells were maintained in fresh Glucose or Galactose medium for 2 hrs. Dashed line represents the threshold for [Fission] beyond which cells were excluded from further data analyses. Metric values were obtained using FL_488_ of MTG probe in the MitoGraph v2.1 analyses. Text describing results in Supplementary results section. **(C)** Box plot showing [Fusion5], [Fission] and [Diameter] in TMRE and MTG stained Cp-resistant A2780 cells identified as ΔΨ^lo^ or ΔΨ^hi^ based on distinct differences in their TMRE uptake. Cells were maintained in fresh Glucose or Galactose medium for 2 hrs. Metric values obtained using FL_488_ of MTG probe in the MitoGraph v2.1 analyses. p values are from non-parametric Kruskal-Wallis test. **(D)** Box plot showing [Fusion5], [Fission] and [Diameter] in Aldh^+^ or Aldh^−^ cells sorted from stained A2780-CP cells. Sorted cells were maintained in fresh RPMI or TIC medium for 24 hrs. Metric values were obtained using FL_633_ of the Mitotracker-633 probe in the MitoGraph v2.1 analyses. p values are from non-parametric Kruskal-Wallis test.

**Figure S4.**
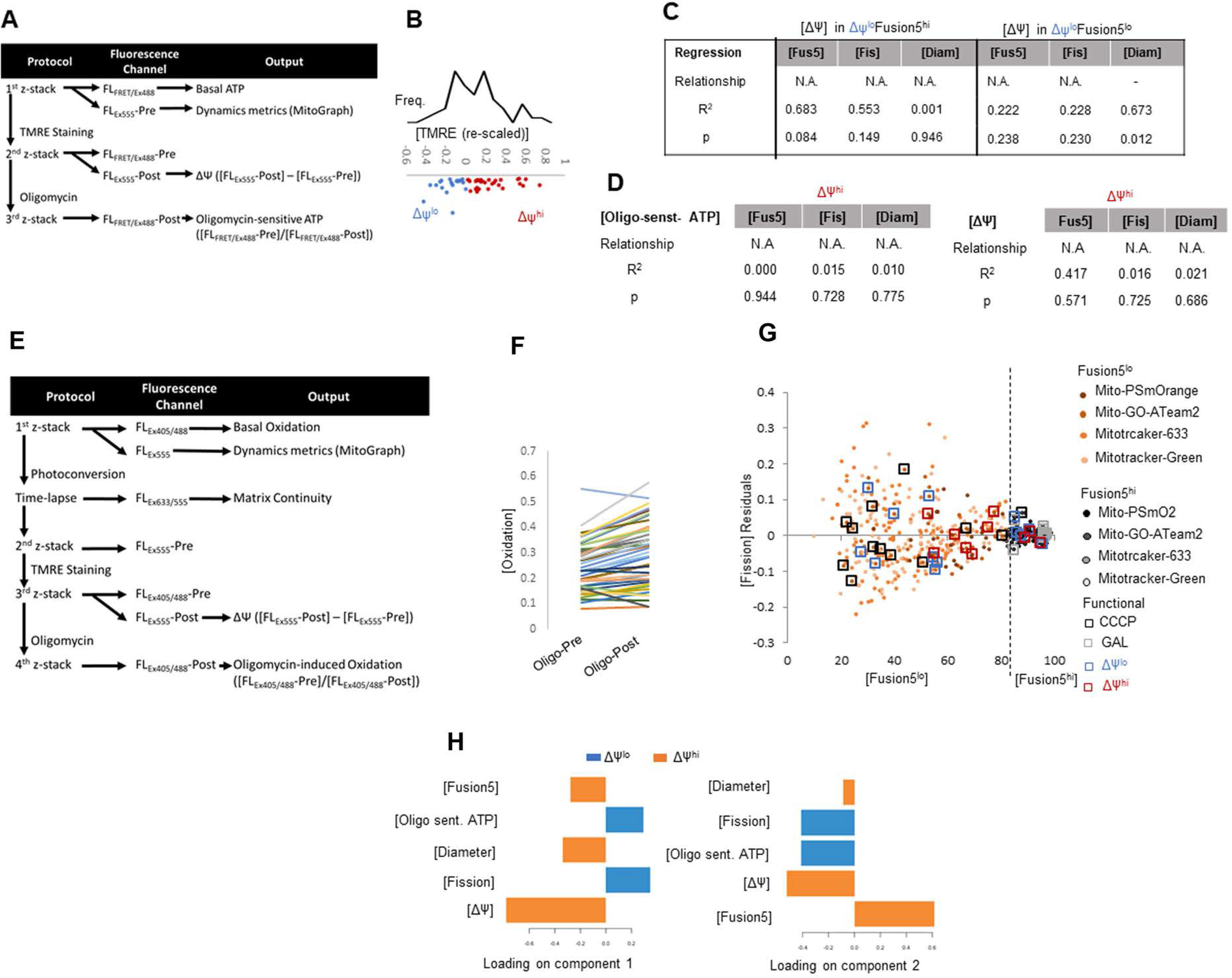
(A) Schematic representing the strategy towards identifying ΔΨ^lo^ and ΔΨ^hi^ cells. The strategy is described in details in Suppl. methods. **(B)** Frequency distribution of ΔΨ^hi/lo^ populations, calculated from their TMRE uptake. Details of the calculation is described in Suppl. methods. **(C)** R^2^, p values and relationship between variables from regression analyses between [ΔΨ] and [dynamics] in Fusion5^hi/lo^ subgroups of ΔΨ^lo^ group. **(D)** R^2^, p values and relationship between variables from regression analyses between [Oligomycin sensitive ATP] (upper panel) or [ΔΨ] (lower panel) and [Fusion5], [Fission] and [Diameter] in the ΔΨ^hi^ population. **(E)** Schematic representing the strategy towards identifying ΔΨ^lo^ and ΔΨ^hi^ cells. The strategy is described in details in Suppl. methods. **(F)** Line plot of change in [Oxidation] with Oligomycin in individual A2780-CP cells. **(G)** Bivariate plot of [Fission] residuals from various previous regression analyses, as mentioned, with respective [Fusion5]; dashed line separates Fusion5^hi/lo^ cells. R^2^ and p values shown. **(H)** Bar plots showing the loading percentages of various mitochondrial metrics on components 1 and 2 of the PLSDA analyses presented in main Figure 6B.

**Figure S5.**
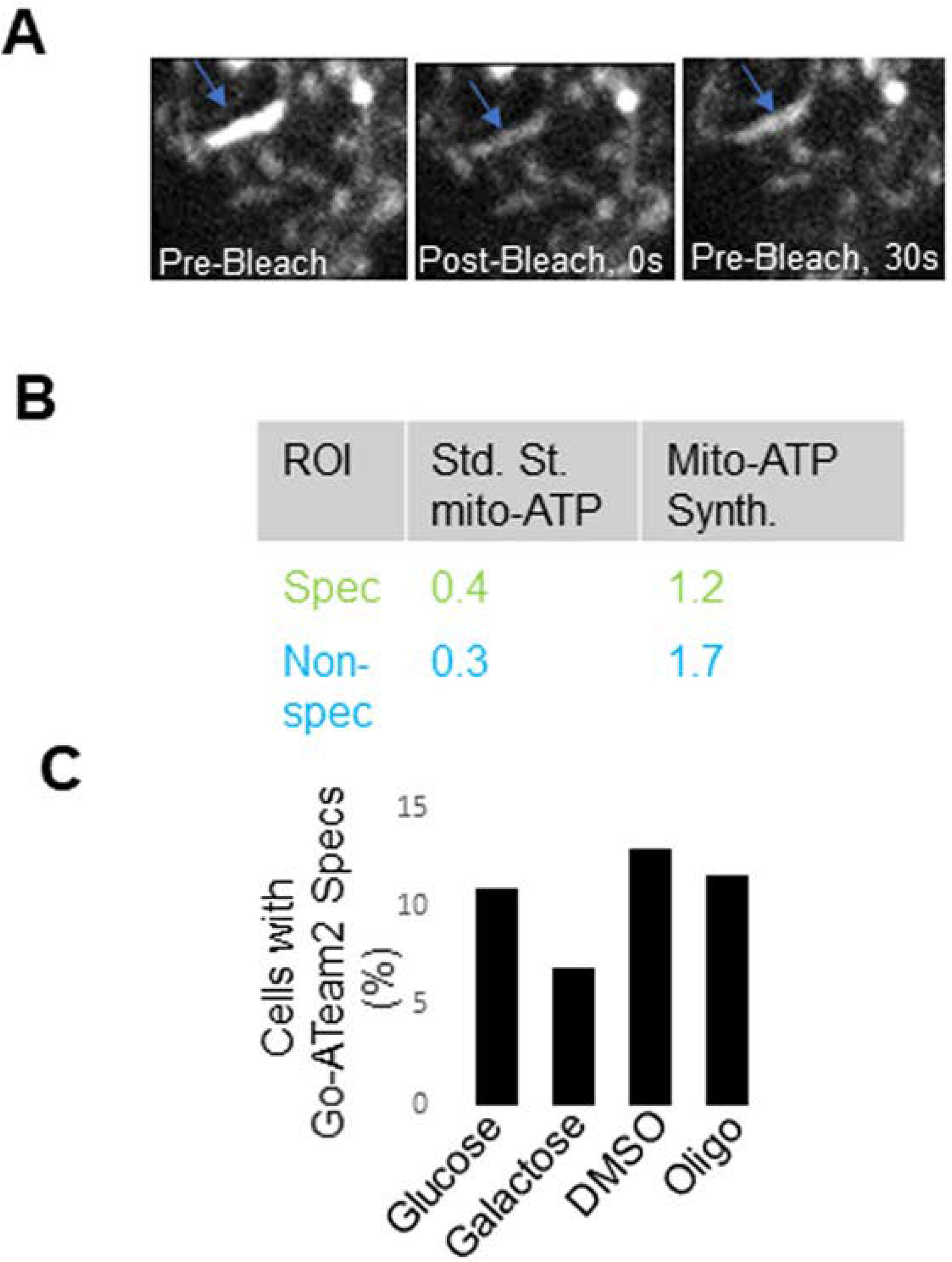
(A) FRAP assay on a mito-GO-ATeam2 enriched mitochondrial speck; arrow on bleaching spot. **(B)** Measurement of FRET from mito-GO-ATeam2 in the speck (blue box in the confocal micrograph) and non-speck (green arrow in the confocal micrograph) regions before and after addition of Oligomycin. FRET values shown in adjacent table. **(C)** Bar plot showing number of the cells with mito-GO-ATeam2 mitochondrial specks in A2780-CP cells in the presence of fresh Glucose, Galactose, DMSO vehicle, or Oligomycin.

